# An expanding arsenal of immune systems that protect bacteria from phages

**DOI:** 10.1101/2022.05.11.491447

**Authors:** Adi Millman, Sarah Melamed, Azita Leavitt, Shany Doron, Aude Bernheim, Jens Hör, Anna Lopatina, Gal Ofir, Dina Hochhauser, Avigail Stokar-Avihail, Nitzan Tal, Saar Sharir, Maya Voichek, Zohar Erez, Jose Lorenzo M. Ferrer, Daniel Dar, Assaf Kacen, Gil Amitai, Rotem Sorek

## Abstract

Bacterial anti-phage defense systems are frequently clustered in microbial genomes, forming defense islands. This genomic property enabled the recent discovery of multiple defense systems based on their genomic co-localization with known systems, but the full arsenal of anti-phage mechanisms in bacteria is still unknown. In this study we report the discovery of 21 new defense systems that protect bacteria from phages, based on computational genomic analyses and phage infection experiments. We find multiple systems with protein domains known to be involved in eukaryotic anti-viral immunity, including ISG15-like proteins, dynamin-like proteins, and SEFIR domains, and show that these domains participate in bacterial defense against phages. Additional systems include protein domains predicted to manipulate DNA and RNA molecules, as well as multiple toxin-antitoxin systems shown here to function in anti-phage defense. The systems we discovered are widely distributed in bacterial and archaeal genomes, and in some bacteria form a considerable fraction of the immune arsenal. Our data substantially expand the known inventory of defense systems utilized by bacteria to counteract phage infection.

## Introduction

To alleviate the burden of phage infection, bacteria evolved a battery of anti-phage defense systems that can collectively be termed the “immune system” of bacteria (Bernheim and Sorek, 2020). Historically, research on bacterial defense systems initially focused on restriction-modification (RM) and abortive infection systems (Abi), and later on CRISPR-Cas (Hampton et al., 2020; Ofir and Sorek, 2018). However, recent studies have shown that the defense arsenal encoded by the prokaryotic pan-genome is much richer than originally anticipated (Doron et al., 2018; Gao et al., 2020). These studies relied on the tendency of defense systems to co-localize on bacterial and archaeal genomes, forming so called “defense islands” (Makarova et al., 2011). Systematic analyses of defense islands in tens of thousands of microbial genomes (Doron et al., 2018; Gao et al., 2020; Rousset et al., 2022) have led to the discovery of several dozens of new defense systems exhibiting a variety of defensive mechanisms. These include systems that utilize second messenger signaling to mediate defense (Cohen et al., 2019; Ofir et al., 2021; Tal et al., 2021a; Whiteley et al., 2019), systems that produce antiviral molecules (Bernheim et al., 2021; Kever et al., 2021; Kronheim et al., 2018), and systems that rely on reverse transcription of small RNAs as part of the defensive machinery (Gao et al., 2020; Millman et al., 2020a).

Despite this substantial progress in mapping the bacterial defense arsenal, it has been estimated that a large number of defense systems have yet to be discovered. A recent study have clustered defense-island-residing genes based on sequence homology, and identified over 7000 gene clusters predicted as encoding new defense genes (Gao et al., 2020). As only a minority of these clusters were tested and characterized experimentally, it is clear that a large diversity of prokaryotic defense systems remain undiscovered.

In this study we set out to further expand the known arsenal of defense systems used by bacteria and archaea to battle phages. By examining clusters of genes enriched in defense islands, we verified previously described defense systems and further discovered 21 new systems. Some of these systems show homology to human and plant genes involved in antiviral innate immune functions, implying that these functions may have evolved from prokaryotic phage-defense systems. Our data show that defense systems can occupy a large proportion of the microbial genome, with some genomes dedicating over one hundred genes for defense against phages.

## Results

To classify proteins into families in a manner not relying on external domain annotation, we clustered ∼135 million bacterial and archaeal proteins from >38,000 microbial genomes into fine-grained clusters based on sequence homology. Each cluster was then analyzed for the propensity of its genes to co-localize with known defense genes in the analyzed genomes. Clusters whose genes showed statistically significant enrichment next to known defense genes (at least 40% of genes co-localizing in defense islands and q value <0.05, see Methods) were considered as possibly functioning in anti-phage defense. Overall we detected 15,574 clusters that passed these thresholds, 24% of which were already known as defensive (Table S1). We then analyzed the genomic neighborhood of genes in each of the defense-associated clusters, to identify cases in which the genes of the cluster are part of a multigene system (Doron et al., 2018).

Due to the multitude of candidate defense systems emerging from this analysis, we prioritized candidates for experimental verification based on their abundance within microbial genomes, the novelty of their predicted protein domains as compared to domains existing in known defense systems, and the availability of the system within a genome of a species closely related to one of our experimental model bacteria, *Bacillus subtilis* and *Escherichia coli*. Overall, we selected 45 candidate defense systems for further analysis. At least two instances for each of these systems, from two different source genomes (when possible) were taken for experimental verification (Table S2). The DNA sequence of each system, spanning the predicted defense genes and the intergenic spaces to include the native promoters (with one exception, see below), was synthesized and transformed into both *E. coli* and *B. subtilis*. These bacteria were then challenged with an array of 26 phages, 12 infecting *E. coli* and 14 infecting *B. subtilis*, spanning a wide range of phage families and life cycles (Table S3). About half of the tested systems, 21 out of 45, protected bacteria against at least one phage in the set, as measured via plaque assay experiments (Figure S1).

### Microbial defense systems with homology to human innate immunity genes

A subset of the microbial defense systems that we identified show homology to proteins involved in the antiviral response of the cell-autonomous innate immune system in animals. These include proteins resembling interferon-stimulated gene 15 (ISG15), dynamin-like proteins, and SEFIR domain proteins. We focused on these systems as they may point to evolutionary connections between antiviral mechanisms in human and bacterial immunity.

#### Homologs of ISG15, a ubiquitin-like protein

The human response to viral infection involves interferon-mediated upregulation of hundreds of antiviral genes (Mesev et al., 2019). Interferon stimulated gene 15 (ISG15) is one of the most strongly and rapidly induced genes as part of the type I interferon antiviral response (Perng and Lenschow, 2018). ISG15 is a ubiquitin-like protein, comprised of two fused ubiquitin-like domains (Narasimhan et al., 2005). This protein was shown to be conjugated by dedicated interferon-induced ubiquitin-conjugating enzymes (E1, E2 and E3) onto many viral and host proteins during infection (Dastur et al., 2006; Dzimianski et al., 2019; Zhao et al., 2004). However, although ISG15 has been known for several decades to protect animal cells against multiple viruses (D’Cunha et al., 1996; Freitas et al., 2020), the exact mechanism of protection is currently unclear (Freitas et al., 2020).

We discovered a 4-gene defense system that comprises a homolog of ISG15, as well as genes with homology to ubiquitin-conjugating enzymes E1 and E2 (Figure 1A; Figure S2). A fourth gene in this operon has homology to ubiquitin-removing enzymes of the JAB/JAMM family (Figure 1A). The bacterial ISG15-like gene encodes, similar to the human ISG15, a protein with two consecutive ubiquitin-like domains, each with clear structural homology to the ISG15 ubiquitin-like domains (Figure S2). The four-gene operon was detected in 110 of the bacterial genomes we studied, and was specifically abundant in Alphaproteobacteria (Table S4). As we could not find a homolog of this system in organisms phylogenetically similar to our lab model organisms, we cloned the system from several different Alpha- and Betaproteobacterial species under an inducible promoter instead of using their native promoters. *E. coli* expressing the prokaryotic ISG15-like system from either *Collimonas, Caulobacter, Cupriavidus, Paraburkholderia*, or *Thiomonas* was broadly protected against infection by multiple phages (Figure 1B). We named genes within this new defense system *bilABCD*, acronym for Bacterial ISG15-Like genes (Figure 1A).

**Figure 1.**
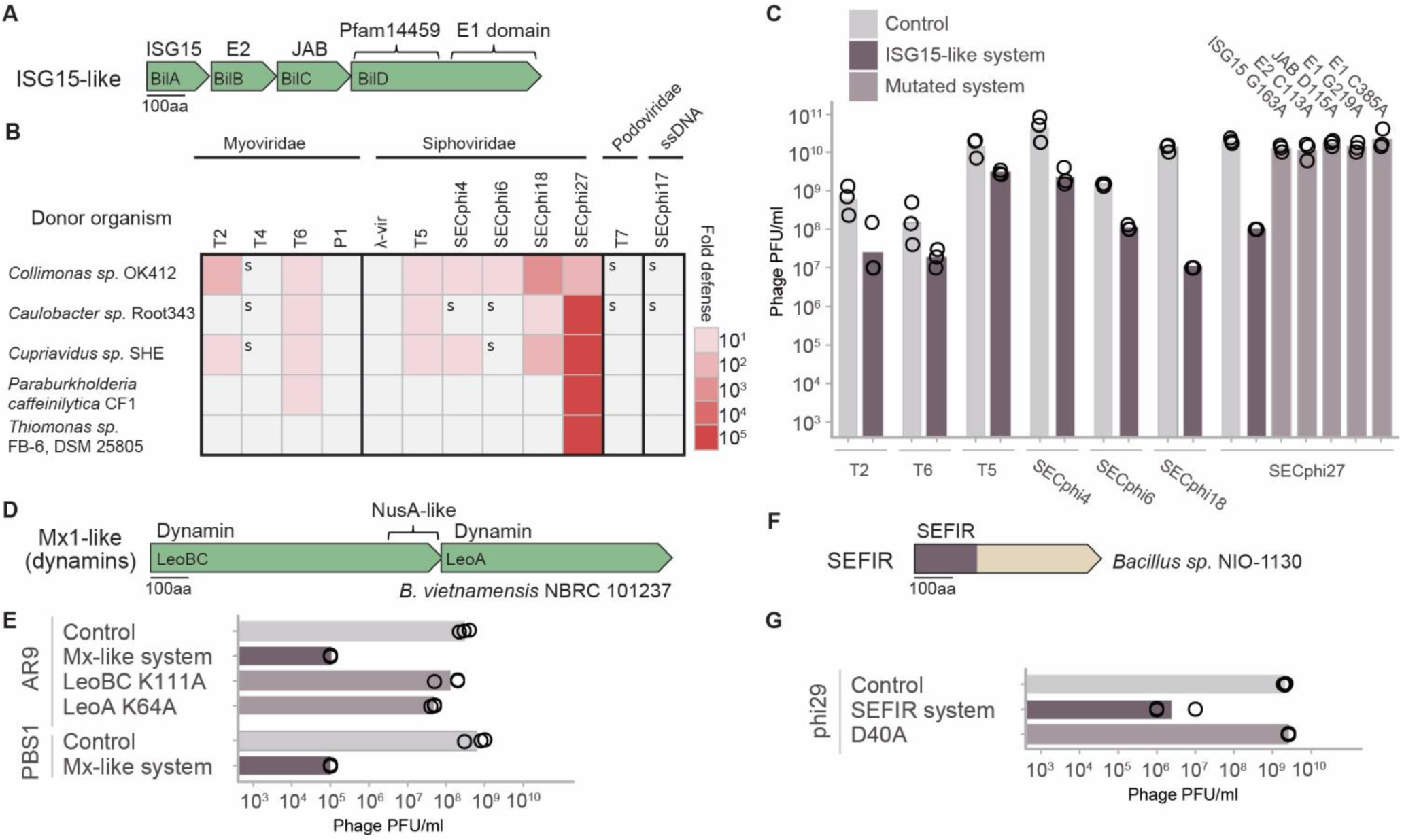
Defense systems encoding domains that participate in human immunity. (A) The ISG15-like operon. (B) Defense profiles of five ISG15-like operons transformed into *E. coli* under an inducible promoter. Fold defense was measured using serial dilution plaque assays, comparing the efficiency of plating (EOP) of phages on the system-containing strain to the EOP on a control strain that lacks the system. Data represent an average of three replicates (See Figure S1). A designation of ‘s’ stands for a marked reduction in plaque size. (C) EOP of phages infecting *E. coli* with and without WT and mutated ISG15-like system from *Collimonas*. Data shown only for phages for which defense was detected. Data represent plaque-forming units (PFU) per ml; bar graph represents an average of three replicates, with individual data points overlaid. (D) A two-gene system with dynamin-like protein domains. (E) EOP of two phages on *B. subtilis* harboring a WT or mutated dynamin-like system from *B. vietnamensis* under native promoter. (F) SEFIR-domain protein from *Bacillus sp*. NIO-1130. The SEFIR domain location is marked on the gene. (G) EOP of phi29 infecting *B. subtilis* harboring the WT or mutant SEFIR from *Bacillus sp*. NIO-1130 under native promoter. EOP data appearing in this figure is also shown in Figure S1.

A point mutation in the conserved glycine residue at the very C-terminus of the ISG15 homolog (G163A), which, in animals, was found to be essential for conjugating the ISG15 onto target proteins, completely abolished defense (Figure 1C). In addition, a point mutation in a conserved cysteine in the predicted active site of the E1 domain (C385A, predicted to inactivate ISG15 binding), as well as a point mutation in the predicted adenylation site of the E1 (G219A) completely impaired defense against phages (Figure 1C). A single amino acid exchange in the predicted active site of the JAB-domain-containing protein also impaired defense (Figure 1C). These results imply that, as in the animal antiviral ISG15, the conjugation of the bacterial ISG15 onto target proteins may be essential for the defensive activity. At this point, the antiviral mechanism of action remains unclear for both the animal and the bacterial ISG15; however, we posit that understanding the mechanism may prove more straightforward in bacteria due to the simplicity of bacteria-phage experimental systems.

The ISG15-like system can be found in a variety of subtypes in microbial genomes. In addition to the operon described above, some systems appear with a single ubiquitin-like domain (instead of the two fused domains in the ISG15-like system above), and in some cases the gene with the JAB deubiquitinase domain is missing (Table S4). Overall such systems are found in 2.6% of the genomes we studied (Table S4). Intriguingly, type II CBASS anti-phage defense systems also contain ancillary genes (*cap2* and *cap3*), which encode enzymes with E1, E2, and JAB domains (Millman et al., 2020b). These enzymes were recently shown to conjugate the cGAS-like defense protein of CBASS onto unknown targets in the bacterial cell to boost immunity (Ledvina et al., 2022). It is possible that the parallel enzymes in the ISG15-like system similarly function to conjugate the ISG15-like protein onto cellular targets as part of the immune function.

#### Dynamin domains, homologs of animal Mx antiviral genes

Dynamins are a family of large GTPases that carry out various membrane remodeling activities, including fission and fusion of vesicles and other subcellular compartments (Ramachandran and Schmid, 2018). A subfamily of eukaryotic dynamins, the Mx proteins, are interferon-induced immune proteins (Haller et al., 2015) that protect against viruses such as influenza, herpes viruses, and members of the bunyavirus family (Crameri et al., 2018; Frese et al., 1996; Krug et al., 1985). It was suggested that Mx GTPases detect viral infection by sensing nucleocapsid-like structures, then trapping viral components and moving them to locations where they become unavailable for the generation of new virus particles (Haller et al., 2015). It was also suggested that Mx proteins actively disassemble viral capsids (Serrero et al., 2022). However, the mechanism of dynamin-like Mx proteins in animal antiviral immunity is not entirely clear.

A recent study showed that a dynamin-family gene called *dynA*, a housekeeping gene in *B. subtilis*, protects against phage infection by maintaining membrane integrity and thus delaying cell lysis (Guo et al., 2022). In our study, we detected an operon of two consecutive dynamin-like proteins that was consistently present next to defense systems in bacterial defense islands (Figure 1D). This operon showed homology to the dynamin-like genes in the *Leo* operon from *Escherichia coli* ETEC H10407 (Michie et al., 2014), which were shown through X-ray crystallography to have dynamin-like structures (Michie et al., 2014) and were suggested to be involved in bacterial virulence due to their localization in a pathogenicity island (Fleckenstein et al., 2000). However, based on the high frequency of co-localization of this operon with known defense systems (60% of the cases), we hypothesized that it has a role in defense against phages. We cloned the two-gene system from *Bacillus vietnamensis* NBRC 101237 into both *B. subtilis* and *E. coli*, and tested whether it had anti-phage activity against the 26 phages in our set. The operon conferred substantial levels of defense, but only against the *B. subtilis* phages AR9 and PBS1, both of which are “jumbo-phages”, encoding unusually large phage particles with genome sizes of >250kb (Figure 1E). Mutating the phosphate binding motif in the GTPase domains of either of the two proteins abolished the defense, suggesting that GTP hydrolysis and the mechanical force it is hypothesized to generate within the bacterial dynamin proteins are necessary for anti-phage defense (Figure 1E).

The two-gene dynamin system we characterized is found in ∼1% of bacterial genomes in our set, belonging to phylogenetically diverse bacterial phyla, including Proteobacteria, Firmicutes, Actinobacteria, Cyanobacteria, Bacteroidetes, and more (Table S5). In some cases, for example in the *E. coli* ETEC H10407 genome, the first gene is divided into two separate genes (Table S5). Although the mechanism of defense of this operon is currently unknown, it is tempting to speculate that it would function similarly to Mx proteins in the animal immune system.

#### Proteins with SEFIR domain

The SEFIR domain is an intracellular signaling domain found in interleukin-17 receptors (IL-17R) and other human immune proteins (Novatchkova et al., 2003; Zhang et al., 2014). Following IL-17 binding, the SEFIR domain of the membrane-bound IL-17R interacts with a SEFIR domain of the cytosolic immune adaptor protein NF-κB activator 1 (Act1), leading to a signaling cascade that activates the immune response (Onishi et al., 2010; Ryzhakov et al., 2011). The SEFIR domain shares structural similarity with Toll/IL-1 receptor (TIR) domains, but structural elements unique to SEFIR and distinct from TIR domains have been described (Zhang et al., 2014).

SEFIR domains were also identified in prokaryotes (Wu et al., 2012; Yang et al., 2018). A crystal structure of a SEFIR domain from *Bacillus cereus* showed high structural similarity to the human IL-17R SEFIR (Yang et al., 2018), and it was hypothesized that bacterial SEFIR domains function as general protein-protein interaction domains for diverse physiological processes (Yang et al., 2018). However, we found that bacterial proteins with SEFIR domains are enriched in defense islands, which suggested a specific role in bacterial immunity. Indeed, a SEFIR-domain protein from *Bacillus sp*. NIO-1130 conferred defense against phage phi29 when expressed in *B. subtilis* (Figure 1F-G). The defense phenotype of the protein was abolished when the amino acid predicted as involved in SEFIR-SEFIR interactions (D40A) was mutated, suggesting that such interactions are necessary for anti-phage defense (Yang et al., 2018). We detected homologs of the SEFIR-domain gene in 460 genomes in our set, belonging to diverse bacteria spanning over 20 phyla, including homologs in the archaeal species *Methanosarcina barkeri* and *Methanosarcina mazei* (Table S6). It is currently unclear whether the eukaryotic SEFIR domain has evolved from the prokaryotic one, or whether the SEFIR domain evolved independently in both taxonomical groups, presumably from TIR domains that have abundant immune functions in both prokaryotes and eukaryotes (Wu et al., 2012).

### A diverse family of Lamassu-like systems involving SMC proteins

Our analysis retrieved multiple operons encoding a protein of the SMC family (Figure 2A). SMC (structural maintenance of chromosomes) is a split ATPase domain in which the two half-ATPase segments are separated by a long coiled coil (Haering et al., 2002). SMC proteins were shown to function in high-order chromosome organization in both prokaryotes and eukaryotes, including chromosome condensation during replication (Michaelis et al., 1997), sister chromatid attachment (Losada et al., 1998), and DNA repair (Potts et al., 2006). The SMC domain was also previously described in a two-gene defense system called Lamassu (Doron et al., 2018). In addition to the SMC gene (*lmuB*), Lamassu encodes the LmuA protein that has an N-terminal Cap4 dsDNA endonuclease domain (previously called DUF4297) (Lowey et al., 2020). Type II Lamassu systems also encode *lmuC*, an additional short gene of unknown function (Payne et al., 2021). It was recently shown that type II Lamassu (called DdmABC system in *V. cholerae*) protects *V. cholerae* from both phage infection and plasmid replication, in a mechanism that involves abortive infection (Jaskólska et al., 2022).

**Figure 2.**
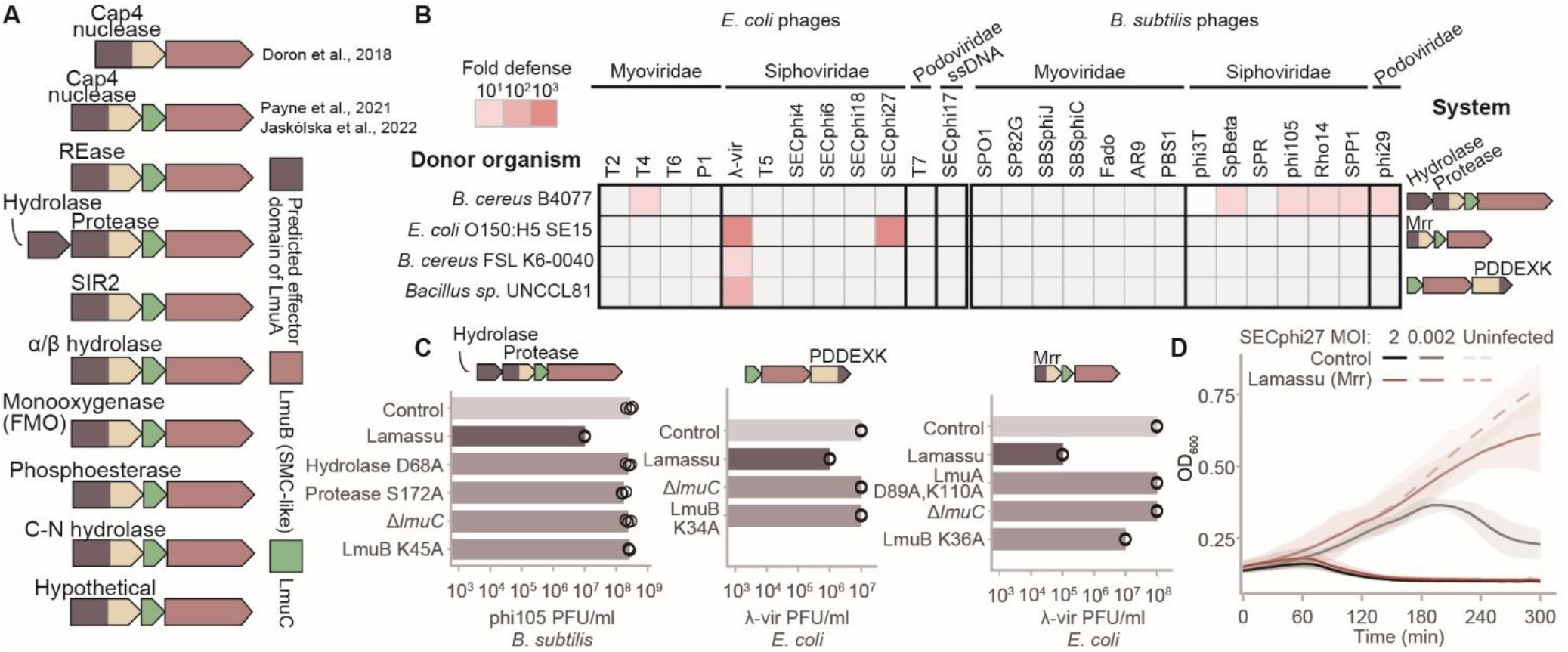
The Lamassu family of defense systems. (A) Domain organization of systems from the Lamassu family. Effector domain at the N-terminus of LmuA is in dark purple. REase stands for restriction endonuclease. (B) Defense profiles of four Lamassu-like operons transformed into *E. coli* and *B. subtilis* under their native promoters. Depiction of fold defense data is as in Figure 1B. (C) EOP of phages infecting WT and mutated Lamassu-like systems. EOP data appearing in this figure is also shown in Figure S1. (D) Growth curves for *E. coli* MG1655 cells harboring a plasmid with the Lamassu-like system cloned from *E. coli* O150:H5 SE15, or negative controls cells containing an empty vector, infected with phage SECphi27 at Time=0. Each growth curve represents the mean of 3 biological replicates, each with an average of 2 technical replicates, and the shaded area corresponds to the 95% confidence interval.

In the current study, we found that the original type I and type II Lamassu systems are only a subset of a large family of Lamassu-like defense systems that are commonly found in bacterial and archaeal genomes (Table S7). Additional members in this family all encode the *lmuB* SMC gene, and most of them also encode the *lmuC* gene. However, the endonuclease domain in *lmuA* can be replaced by other domains, including endonuclease domains of the PDDEXK and Mrr families, sirtuin (SIR2) domain, and additional hydrolase, protease, and monooxygenase domains (Figure 2A). Many of these domains function as effectors in other bacterial defense systems that protect through abortive infection. For example, the Cap4 endonuclease functions as the effector domain in CBASS systems, where it cleaves both host and phage DNA when activated (Lowey et al., 2020). Moreover, SIR2 domains cause NAD^+^ depletion in many abortive infection systems, including Thoeris, DSR, and more (Garb et al., 2021; Ofir et al., 2021; Zaremba et al., 2021). It is therefore likely that systems in the Lamassu family all protect against phage via abortive infection.

We selected 6 instances of the expanded Lamassu family to be tested experimentally. Of these, four systems showed defense against *E. coli* and/or *B. subtilis* phages (Figure 2B). In all cases, defense was dependent on a functional SMC ATPase enzyme, because a point mutation in the ATPase active site of the SMC gene abolished defense (Figure 2C). Mutation in the effector domains of these systems also abolished defense, as well as the deletion of *lmuC*, demonstrating that all components are necessary for the defensive function (Figure 2C). Infection in liquid culture showed that Lamassu-like systems protect the culture when phages are supplied in low multiplicity of infection (MOI), but the culture collapses when phages are added in high MOI (Figure 2D). This phenotype is the hallmark of abortive infection systems (Lopatina et al., 2020), confirming that other Lamassu-like systems, in addition to the DdmABC type II Lamassu (Jaskólska et al., 2022), also function via abortive infection.

Systems from the expanded family of Lamassu are found in ∼10% of the bacterial and archaeal genomes in our set (Table S7). Members of this family were previously detected in regions within prophage genomes that form hotspots for bacterial immune systems (Rousset et al., 2022), further supporting a general role in defense. As the *lmuA* gene frequently encodes an effector domain that is known to execute abortive infection in other defense systems, we hypothesize that the SMC-containing LmuB protein is responsible for the recognition of the invading phage, as was also previously suggested in other studies (Jaskólska et al., 2022; Krishnan et al., 2020). As SMC proteins typically interact with DNA, it is possible that LmuB primarily detects the DNA of the invading phage, possibly by recognizing phage DNA replication intermediates, as also proposed by a recent study (Jaskólska et al., 2022). Following phage DNA recognition, ATP hydrolysis by the LmuB ATPase would activate LmuA, via a mechanism that can be similar to the SMC-containing DNA repair system SbcCD, in which recognition of damaged DNA activates an associated nuclease (Käshammer et al., 2019).

### Multiple additional defense systems

In addition to the systems we described above, multiple other systems were discovered here (Figure 3; Tables S8-S24). We named many of these systems after protective deities from various world mythologies, as previously done for newly discovered anti-phage systems (Doron et al., 2018). The set of discovered systems includes genes spanning a variety of protein domains and molecular functions (Figure 3; Tables S8-S24). For example, we found a family of systems, that we call here Mokosh (Slavic goddess protector of women’s destiny), which comprises genes with an RNA helicase domain of the COG1112 family and a predicted nuclease of the PLD family (Figure 3A). The RNA helicase domain of the Mokosh system shows substantial homology to the human RNA helicase Upf1, which regulates degradation of RNA transcripts that contain premature stop codons (Figure S3) (Kim et al., 2005). In type I Mokosh, the RNA helicase and PLD nuclease are separated into two genes, and the gene with the helicase domain additionally includes a serine-threonine kinase domain (STK) (Figure 3A). Type II Mokosh systems comprise a single gene with the RNA helicase at the N-terminus and the PLD nuclease at the C-terminus (Figure 3A). Three Mokosh systems in our set showed defense against phages in *E. coli* (Figure 3B). Mutations in the ATP-binding domain of the helicase abolished defense, as well as mutations in the active site of the PLD nuclease (Figure 4). We hypothesize that this system involves recognition of phage RNA, or RNA/DNA intermediates associated with phage replication.

**Figure 3.**
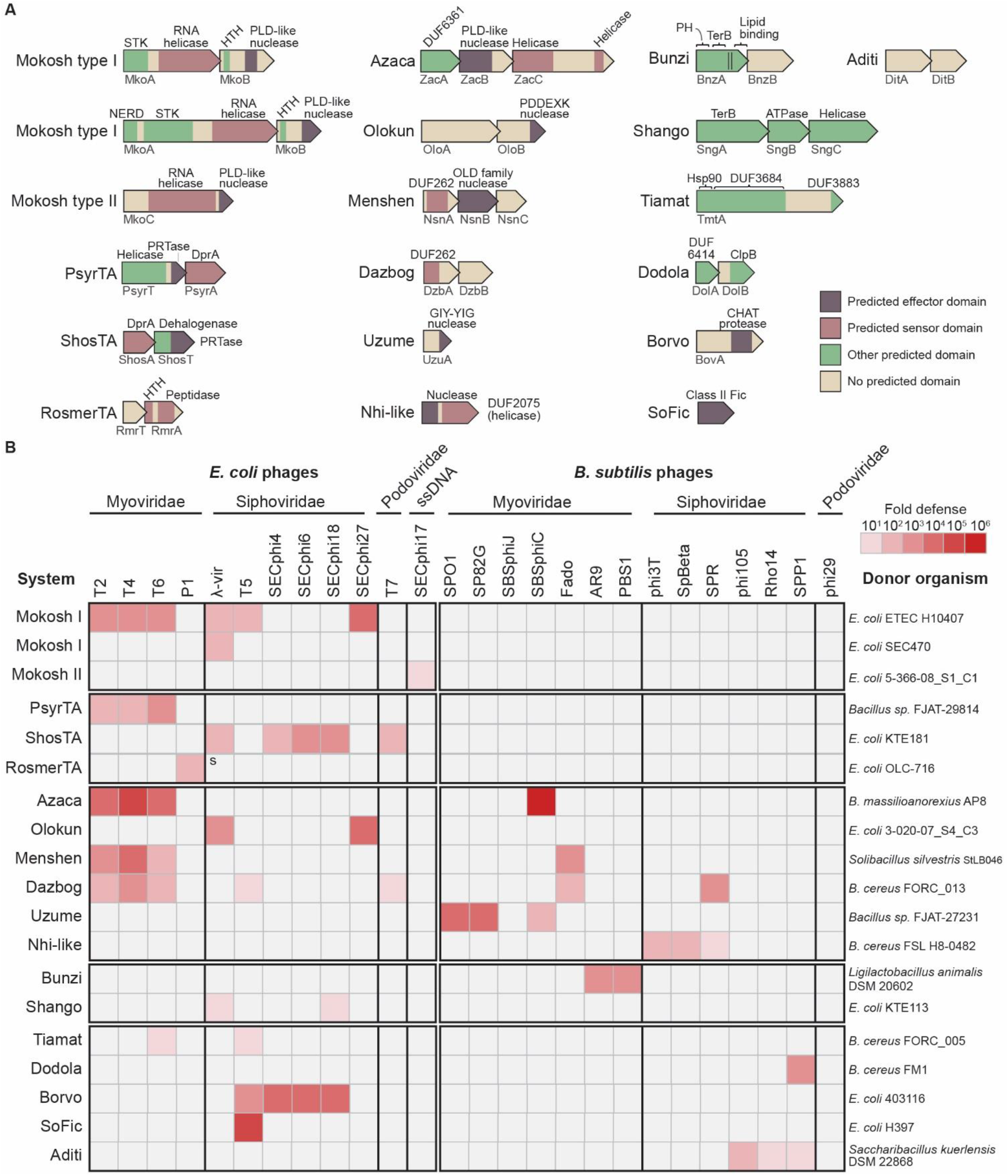
Additional defense systems discovered in this study. (A) Domain organization of defense systems. The two lines of BnzA represent the predicted transmembrane helices. (B) Defense profiles of defense systems operons transformed into *E. coli* and *B. subtilis* under their native promoters. Depiction of fold defense data is as in Figure 1B, source data is shown in Figure S1.

**Figure 4.**
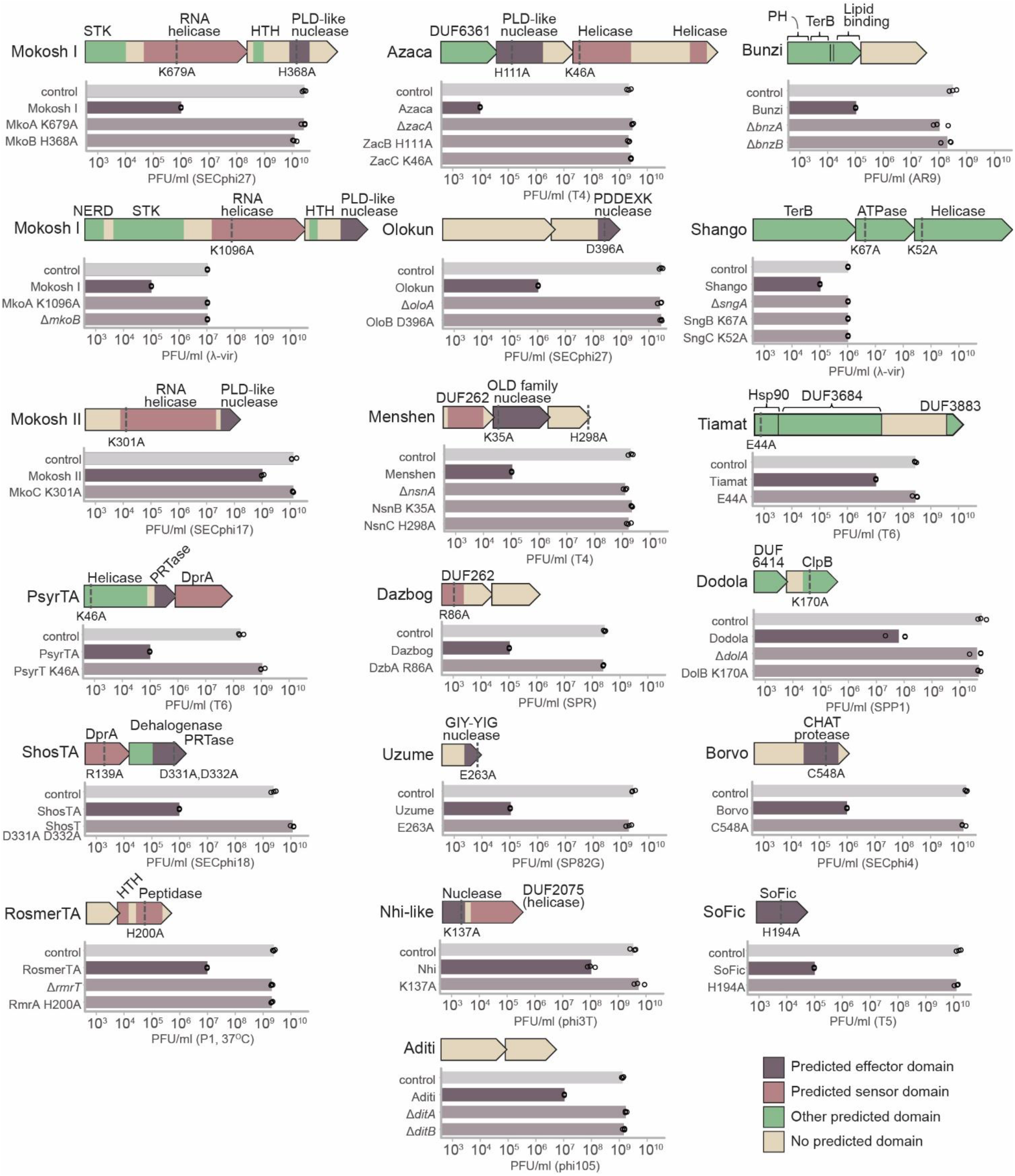
EOP of phages infecting WT and mutated systems. EOP data appearing in this figure is also shown in Figure S1. All experiments were done at room temperature except for the RosmerTA system, for which experiments were performed at 37°C. Donor organisms are as mentioned in Figure 3B. We were unable to transform the following deletion mutants, possibly due to toxicity: Δ*psyrA*, Δ*shosA*, Δ*rmrA*, Δ*dzbB*.

Several two-gene systems that we identified here as involved in phage resistance were previously shown to encode toxin antitoxin (TA) systems (Figures 3-4). These include the ShosTA system (Kimelman et al., 2012), that was also recently verified as phage-defensive in another study (Rousset et al., 2022), and the PsyrTA system (also called RqlHI) (Russell and Mulvey, 2015; Sberro et al., 2013). Both these systems encode an antitoxin that has homology to DprA, a single-stranded DNA (ssDNA)-binding protein known to be involved in DNA transformation (Mortier-Barrière et al., 2007). The toxin protein contains a phosphoribosyl transferase (PRTase) domain, which was previously found in effectors of retron abortive infection systems (Mestre et al., 2020; Millman et al., 2020a). Another defense system in our set, which we call here RosmerTA (Gallo-Roman goddess of fertility) involves an antitoxin with a Zn-peptidase and a toxin of unknown function, a configuration previously shown in TA systems (Luo et al., 2015; Matelska et al., 2017; Rousset et al., 2021). These systems join other TA systems, including RnlAB (Koga et al., 2011), ToxIN (Blower et al., 2011), and DarTG (LeRoux et al., 2021), which were shown to confer phage defense (LeRoux and Laub, 2022).

Two systems in our set, which we denote Bunzi and Shango (Kongo goddess of rain and Yoruba god of thunder), encode proteins with TerB domains (Figure 3-4). TerB is a protein that was originally identified in the tellurite resistance gene cassette (Whelan et al., 1997), and was hypothesized to be involved in phage defense (Anantharaman et al., 2012). Although the exact function of this domain is unknown, it was shown to be associated with the periplasmic membrane (Alekhina et al., 2011), and indeed the TerB protein in the Bunzi system contains predicted transmembrane-spanning helices as well as a domain predicted to interact with lipids (Figure 3A). Possibly, these systems may be involved in membrane surveillance. Notably, the Shango system has also been discovered in a parallel study, where it was shown to confer anti-phage defense in *Pseudomonas aeruginosa* (J. Bondy-Denomy and A. Davidson, personal communication).

Additional protein domains found in the defense systems described here include various nuclease, peptidase, and chaperon-associated domains. We also found a standalone protein with Fic domain (SoFic), which is known to be a protein AMPylase (capable of ligating AMP onto proteins) (Engel et al., 2012; Harms et al., 2016). One defensive protein in our set encodes a predicted N-terminal nuclease and the domain of unknown function DUF2075 (pfam: PF09848), a domain organization identical to the Nhi anti-phage protein discovered recently in *Staphylococcus epidermidis* (Bari et al., 2022). Although there is no sequence identity between the *S. epidermidis* Nhi and the protein we studied here, we denoted our protein Nhi-like, because the similar domain organization suggests a similar function. One defense system, which we call here Aditi (Hindu guardian goddess of all life), contains two proteins of unknown function, which had no homology to any known domain even when using the sensitive homology prediction tool HHpred (Steinegger et al., 2019; Zimmermann et al., 2018).

### Distribution of defense systems in microbial genomes

To assess the distribution of defense systems in prokaryotic genomes, we updated DefenseFinder (Tesson et al., 2021) with all of the systems described in this study and applied it on a subset of 3,895 genomes in our database that were defined as “finished”, i.e., fully sequenced and assembled with no gaps. In total we found 22,610 defense systems in 3,688 of the 3,895 genomes (Figure 5A, Tables S25, S26). Of these, 12.2% are systems described in this study (Figure 5B). Of the 3,895 genomes analyzed, 1,837 (47%) contain at least one of the systems described in this study (Table S25).

**Figure 5.**
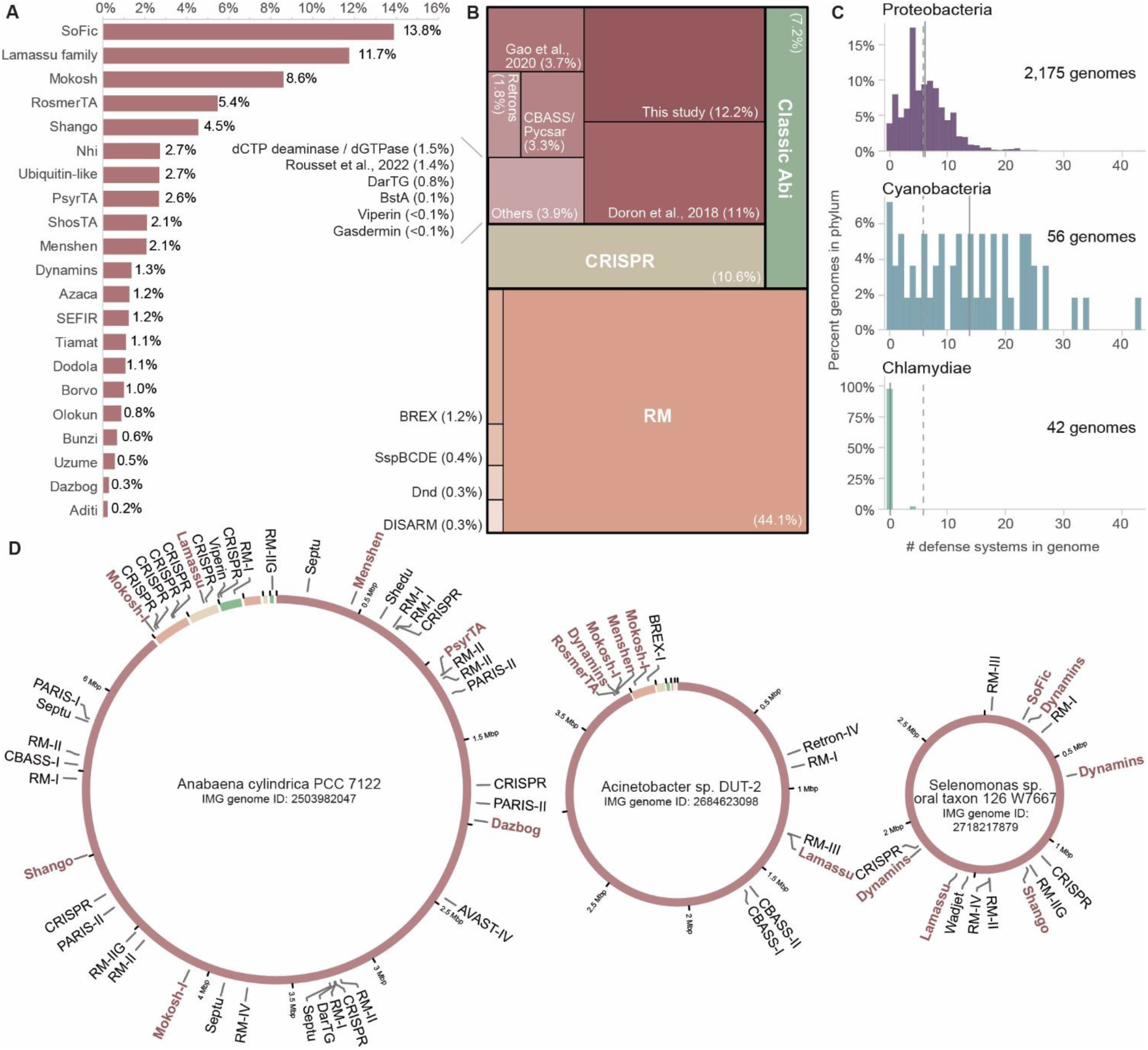
Distribution of defense systems in microbial genomes. (A) Presence of defense systems described in this study in finished genomes. Y-axis depicts the fraction of genomes, out of the 3,895 finished genomes analyzed, that contain each of the defense systems discovered in this study. (B) Abundance of defense systems in finished genomes. Fraction out of 22,610 systems found in 3,895 finished genomes. (C) Phylum-specific distribution of defense systems for selected phyla. X-axis presents the number of defense systems per genome; Y-axis presents the fraction of genomes within the phylum. Dashed line is the average number of defense systems per genome of the entire set of 3,895 genomes; Solid gray line is the average within the phylum. Figure S4 presents data for additional phyla. (D) Presence of defense systems in representative genomes. Colors in the ring represent different genomic scaffolds. Names of defense systems detected in this study are in brown.

In line with previous reports (Makarova et al., 2020; Millman et al., 2020b; Oliveira et al., 2014; Tal et al., 2021b; Tesson et al., 2021) the most common systems are restriction modification systems (found in 84% of the genomes we analyzed), CRISPR (45% of the genomes), and CBASS/Pycsar (17% of the genomes) (Table S25). We found that bacteria and archaea hold on average 5.8 defense systems per genome, and dedicate on average 15.4 genes for defense. From a phylogenetic perspective there are several phyla that largely deviate from this average. For example, in line with previous reports (Tesson et al., 2021) we found that genomes of the obligate intracellular species Chlamydiae encode no known defense systems, even after including the new systems described in the current study (Figure 5C; Figure S4). On the other hand, Cyanobacterial genomes encode, on average, 13.8 defense systems and the distribution around the average is very wide (between 0 to 43 systems per genome) (Figure 5C). Examining archaeal genomes available in our set, we find that genomes of the phylum Crenarchaeota (24 genomes of 15 different species) rely almost completely on CRISPR as a defense mechanism (2.4 CRISPRs on average), with only 6 genomes encoding an additional RM system, and no other system (Table S25). In contrast, Euryarchaeota species encode a diverse arsenal of systems, with an average of 7.5 systems per genome (Figure S4).

Some genomes encode an exceptionally large number of defense systems. For example, *Anabaena cylindrica* PCC 7122 encodes 13 RM loci, 10 CRISPR loci, and 20 additional defense systems, which are spread in defense islands throughout the genome (Figure 5D). This organism seems to devote at least 132 genes for defense against phage. In some organisms, the new systems we reported in the current study form a substantial fraction of the defensive arsenal (Figure 5D). On the other hand, there are 207 genomes in our dataset (5.3%) in which we could not find any known defense system. Most of these genomes are small (82.6% have <2,000 genes), but this set also includes strains of well-studied free-living bacteria (Table S26). It is possible that these strains encode defense systems that are not yet discovered, or are otherwise especially vulnerable to phage.

## Discussion

Our study substantially expands contemporary knowledge on the defense arsenal used by bacteria to mitigate phage infection. Beginning with the Doron et al. publication in 2018 (Doron et al., 2018), studies that relied on genomic localization signatures unearthed, together with the current study, over 50 previously unknown defense systems (Bernheim et al., 2021; Cohen et al., 2019; Doron et al., 2018; Gao et al., 2020; Millman et al., 2020a; Rousset et al., 2022; Tal and Sorek, 2022; Tal et al., 2021a, 2021b). A minority of these systems were studied in depth, revealing a wide functional variety of defensive mechanisms involving new signaling molecules (Cohen et al., 2019; Ofir et al., 2021; Tal et al., 2021b; Whiteley et al., 2019), chemical defense (Bernheim et al., 2021), reverse transcription of non-coding RNAs (Bobonis et al., 2020; Gao et al., 2020; Millman et al., 2020a), and depletion of essential molecules during infection (Garb et al., 2021; Tal et al., 2021a). It is therefore anticipated that future in-depth studies of the systems described here will reveal similarly diverse mechanisms of defense.

While some of the defense systems we discovered, e.g., the SoFic, Lamassu-like and Mokosh systems, are found in a substantial fraction of microbial genomes, many of the systems discovered here are rare (Figure 5A). The long tail of rare defense systems that sparsely populate bacterial genomes may be the consequence of efficient counter-defense measures developed by phages over the course of evolution, which may have resulted in subsequent elimination of these systems from many of the genomes. Nevertheless, rare defense systems were previously found as useful for biotechnological and biomedical uses. For example, the Cas12 and Cas13 proteins used for molecular diagnostics (Singh et al., 2022), as well as the prokaryotic viperin enzymes that produce candidate antiviral molecules (Bernheim et al., 2021), are all present in less than 1% of the genomes analyzed. Based on the past adoption of bacterial defense systems as molecular tools, it is possible that some of the defense systems we discovered may form the basis for future such tools, even if they are rare in nature.

In recent years it became clear that multiple components of the human innate immune system have evolved from bacterial anti-phage defense mechanisms (Wein and Sorek, 2022). CBASS systems were shown to be ancestral to the human cGAS-STING pathway (Cohen et al., 2019; Millman et al., 2020b; Morehouse et al., 2020; Whiteley et al., 2019; Ye et al., 2020), and evidence for eukaryotic acquisition of the antiviral protein viperin from archaea have been presented (Bernheim et al., 2021). Moreover, homologs of human immunity proteins such as gasdermins, Argonautes, and proteins with TIR domains were all documented in bacteria and were shown to participate in phage defense (Johnson et al., 2022; Koopal et al., 2022; Kuzmenko et al., 2020; Morehouse et al., 2020; Ofir et al., 2021). In the current study, we report on additional bacterial defense systems that encode domains known to be involved in animal immunity. Intriguingly, in some cases, such as the ISG15 and Mx antiviral proteins, the mechanism by which they confer defense in human cells is not entirely clear despite decades of research (Haller and Kochs, 2020). Although their mechanisms of action in bacteria are also unknown at this stage, the relative simplicity of bacteria-phage experimental models may result in more rapid mechanistic solutions for these systems when studied in bacteria. We therefore envision that studies of these defense systems in bacteria may eventually shed light on their parallel roles in animal immunity.

While our study revealed a substantial number of defense systems, many gene families that showed high propensity of localization within defense islands were not studied here due to their rarity in nature and the limited experimental capacity of our study (Table S1). Moreover, some of the tested candidate defense systems that were not verified as defensive in this study may still have defensive roles under conditions not tested in this study, or against types of phages that were not included in our set of 26 phages. Therefore, despite the remarkable progress in understanding the bacterial immune system in the past few years, we are still far from understanding the full capacity of the bacterial immune arsenal.

## Acknowledgements

We thank the Sorek laboratory members for comments on earlier versions of this manuscript. We also thank M. Steinegger and M. Mirdita for assistance with MMseqs2 analyses, and S. Afgin and V. Golodnitsky for assistance with high performance computations. A.M. was supported by a fellowship from the Ariane de Rothschild Women Doctoral Program and, in part, by the Israeli Council for Higher Education via the Weizmann Data Science Research Center. A.B. is supported by a CRI Research Fellowship from the Bettencourt Schueller Foundation, the ATIP-Avenir program from INSERM (R21042KS / RSE22002KSA) and the Emergence program from the University of Paris-Cité (RSFVJ21IDXB6_DANA). J.H. was funded by the Deutsche Forschungsgemeinschaft (DFG, German Research Foundation), grant 466645764. G.O. was supported by the Weizmann Sustainability and Energy Research Initiative (SAERI) doctoral fellowship. M.V. is a Clore Scholar and was supported by the Clore Israel Foundation. R.S. was supported, in part, by the European Research Council (grant ERC-AdG GA 101018520), Israel Science Foundation (grant ISF 296/21), the Deutsche Forschungsgemeinschaft (SPP 2330, grant 464312965), the Ernest and Bonnie Beutler Research Program of Excellence in Genomic Medicine, the Minerva Foundation with funding from the Federal German Ministry for Education and Research, and the Knell Family Center for Microbiology.

## Methods

### Computational prediction of defense systems

#### Generating protein clusters

A database of 38,167 bacterial and archaeal genomes was downloaded from the Integrated Microbial Genomes (IMG) database in October 2017 (Chen et al., 2019). All proteins in the database were first clustered using the ‘cluster’ option of MMseqs2 (release 2-1c7a89) (Steinegger and Söding, 2017), with default parameters (C1 clusters). To remove redundancy, the clusthash option of MMseqs2 was used with --min-seq-id 0.95 and then only one representative of each redundancy group was included. Clusters were further aggregated into larger clusters (C2 clusters) using four additional cycles of clustering, in which - in each cycle - a representative sequence was taken from each cluster using the ‘createsubdb’ option of MMseqs2 and representative sequences were clustered using the ‘cluster’ option with the ‘--add-self-matches’ parameter. For the first additional clustering cycle, the ‘cluster’ option was run with default parameters; for the additional cycles 2–4, clustering was run with sensitivity parameter ‘-s 7.5’, and for the additional cycle 4, the ‘–cluster-mode 1’ parameter was also added.

#### Defense score calculation

The pfam and COG annotations of all proteins in the set were downloaded from IMG in October 2017 (Chen et al., 2019). A set of pfams and COGs of verified defensive protein families was compiled based on Doron et al. (Doron et al., 2018), and all proteins annotated with at least one of these pfams or COGs were considered as known defense proteins (‘positive set’). For each C1 cluster, the fraction of genes associated with known defense genes (‘defense score’) was computed as previously described (Cohen et al., 2019; Doron et al., 2018). To minimize false predictions, positive set proteins whose C1 clusters received a score lower than 0.25 were removed from the positive set and the score calculation was reiterated. Defense score was then calculated for C2 clusters.

To assess whether the defense score is higher than expected by chance, a p-value was calculated for the defense score of each C2 cluster as follows: for each genome-of-origin of each gene in the cluster, the number of genes that had at least one positive set gene in their environment (up to 10 genes upstream or downstream) was counted. These numbers were summed, and divided by the total number of genes present in the genomes-of-origin of the genes in the cluster. The resulting value represents the probability of a random gene within the genome set to be present next to a known defense gene. This value, calculated for each cluster, was then used to calculate the probability to obtain the defense score of the cluster (or higher), assuming a binomial distribution, yielding a p-value. This p-value was corrected for multiple testing using the qvalue package of R (Storey et al., 2019), generating a q-value for each C2 cluster.

#### Prediction of multi-gene defense systems

Multi-gene defense systems were predicted as previously described (Doron et al., 2018). For this, C2 clusters containing genes from at least 4 genera, an effective cluster size >=10, a defense score >=0.4 and q-value <=0.05 were selected as anchor clusters for system prediction (Table S1). Effective size was calculated as the number of genes within the cluster that resided in a long-enough genomic scaffold such that 10 genes from each of their sides were present on the scaffold. To predict the domain organization of the systems, all proteins were scanned with HHpred (Zimmermann et al., 2018) against the PDB, COG, and Pfam databases.

### Experimental validation of defense systems

#### Cloning of candidate defense systems

All sequences and source details for the systems cloned in this study are listed in Table S2. The predicted defense systems were synthetized and cloned in the shuttle vector pSG1-RFP (between the AscI and NotI sites of the multiple cloning sites) by Genscript Corp. as previously described (Doron et al., 2018). The ISG15-like systems were cloned into an arabinose-inducible plasmid (pBAD, Thermo Fisher #V43001). The cloned vector was transformed into *E. coli* MG1655 cells (ATCC 47076) and verified with PCR. The same vector was also transformed into *B. subtilis* BEST7003 cells (kindly provided previously by M. Itaya) using 10X MC medium (80 mM K2HPO_4_, 2% glucose, 30 mM trisodium citrate, 22 μg/ml ferric ammonium citrate, 0.1% casein hydrolysate, 0.2% potassium glutamate, 30 mM KH_2_PO_4_) (Wilson and Bott, 1968). The system was integrated into the *amyE* locus of *B. subtilis*, and resulting transformants were screened on starch plates for amylase-deficient phenotype.

Whole-genome sequencing was then applied to all transformed *B. subtilis* and *E. coli* clones as described in (Goldfarb et al., 2015) to verify system integrity and lack of mutations. As a negative control for systems expressed in *B. subtilis*, a strain with an empty pSG1 plasmid, containing only the spectinomycin-resistance gene in the *amyE* locus, was used. As a negative control for systems expressed in *E. coli*, the wild-type *E. coli* MG1655 carrying an empty pSG1 plasmid was used. As a negative control for systems expressed in *E. coli* on pBAD (ISG15-like systems only), *E. coli* MG1655 carrying pBAD expressing sfGFP was used.

For strains with gene deletions and point mutations, plasmids containing systems with these deletions/mutations were commercially synthesized by Genscript. The mutated systems were transformed into *B. subtilis* and *E. coli* as described above, and clones used were fully sequenced to verify proper integration and sequence of the mutated systems.

#### Phage strains, cultivation, and plaque assay

The phages used in this study and their sources are listed in Table S3. Phage Fado (Genbank: OM236516.1) was isolated in this study from a soil sample on *B. subtilis* BEST7003 as previously described (Doron et al., 2018). To sequence the Fado phage genome, 500 µl of the phage lysate was treated with DNase-I (Merck cat #11284932001) added to a final concentration of 20 µg/ml and incubated at 37C for 1 hour to remove bacterial DNA. Then DNA was extracted using the QIAGEN DNeasy blood and tissue kit (cat #69504) and DNA libraries were prepared using a modified Nextera protocol (Baym et al., 2015) for Illumina sequencing. Phage DNA was assembled from sequenced reads using SPAdes v. 3.11.0 (with the –careful and –cov-cutoff auto modifiers) (Bankevich et al., 2012), genes were annotated using the online RAST server (Aziz et al., 2008) and the sequence was deposited in GenBank under the accession number OM236516.

Phages were propagated on either *E. coli* MG1655 or *B. subtilis* BEST7003 by adding the phage to a bacterial liquid culture grown to OD_600_=0.3 and incubated at 37°C until culture collapse. Propagated phage lysates were centrifuged at 4000 RPM to pellet bacterial cells and the supernatant was filtered through a 0.2 µM filter. Lysate titer was determined using the small drop plaque assay method as previously described (Mazzocco et al., 2009).

To measure defense, plaque assays were performed on bacterial strains expressing the defense system and a negative control strain lacking the system. For this, an overnight culture of bacteria was mixed with pre-melted MMB agar (LB + 0.1 mM MnCl_2_ + 5 mM MgCl_2_ + 5 mM CaCl_2_ + 0.5% agar), and let to dry at room temperature for 1 hour. Then, 10 µl drops from 10-fold serial dilutions of the phage lysate in MMB were dropped on top of the solidified layer containing bacteria. After the drops dried up, plates were incubated overnight at room temperature (or 37°C for the RosmerTA system) and the plaque forming units (PFUs) per ml were counted. Fold defense was calculated as the ratio between the PFU/ml values retrieved using the same phage lysate on control bacteria and bacteria containing the candidate defense system. In the case of ISG15-like systems, expression was induced by the addition of 0.2% arabinose.

#### Phage-infection dynamics in liquid medium of the Lamassu-family systems

Overnight cultures of *E. coli* MG1655 with Lamassu-Mrr or an empty pSG1 plasmid were diluted 1:100 in MMB medium supplemented with ampicillin and incubated at 37°C while shaking at 200 rpm until early log phase (OD_600_ = 0.3). 180 μl of the diluted culture were transferred into wells in a 96-well plate containing 20 μL of phage SECphi27 lysate for a final MOI of 2 or 0.002, or 20 μL of phage buffer (50 mM Tris pH 7.4, 100 mM MgCl2, 10 mM NaCl) for uninfected control. Infections were performed in duplicate from overnight cultures prepared from two separate colonies. Plates were incubated at 25°C with shaking in a TECAN Infinite200 plate reader and OD_600_ was followed with measurement every 10 min.

#### Building DefenseFinder HMM Models

In order to build hidden markov models (HMMs) for usage in DefenseFinder, we first collected, for each gene in each verified system, the set of sequences representing its protein family (Table S4-S24). For multi-gene systems, the C2 clusters and pfams that participate in the system were collected and searched for in all genomes. When all clusters or pfams of a system were found closely localized on a genome (allowing insertions of genes up to double the size of the verified system), this genome was recorded as containing the system.

For single gene systems, all genes of the C2 cluster from which the experimentally verified gene(s) originated, were considered as belonging to the gene family. If there were additional C2 clusters that had a similar domain organization, a defense score of >0.3, and similar average protein size, genes from these clusters were also considered as belonging to the protein family.

For the bacterial SEFIR, due to the similarity to TIR domains, all proteins of the clusters were searched with HHblits (Steinegger et al., 2019) against uniclust30 2018 (Mirdita et al., 2017) and then searched with HHsearch (Steinegger et al., 2019) against the PDB_mmCIF70 (Berman et al., 2000) and Pfam32 (El-Gebali et al., 2019) databases. Proteins where the SEFIR hit was located in the N-terminus and was stronger than the TIR hit were selected as instances of the SEFIR gene.

HMM profiles were built for each gene of each system as described previously (Tesson et al., 2021). Briefly, sequences for each protein were aligned using MAFFT v7.475 (Katoh et al., 2002) (default options, mode auto) and then used to produce protein profiles with Hmmbuild (default options) from the HMMer suite v3.3 (Eddy, 2011). HMM GA score threshold was assigned manually by inspecting the distribution of the scores.

#### Structural alignments

The structures for ISG15-like BilA (IMG ID: 2609810443) and the helicase domain of Mokosh MkoA (IMG ID: 2597791081) were predicted using AlphaFold2 v2.1.1 (Jumper et al., 2021). Best ranking structures were then aligned using PDBeFold (Krissinel and Henrick, 2004) to the corresponding PDB entries (3RT3 and 6EJ5 for BilA and MkoA, respectively).

## Supplementary figures

**Figure S1.**
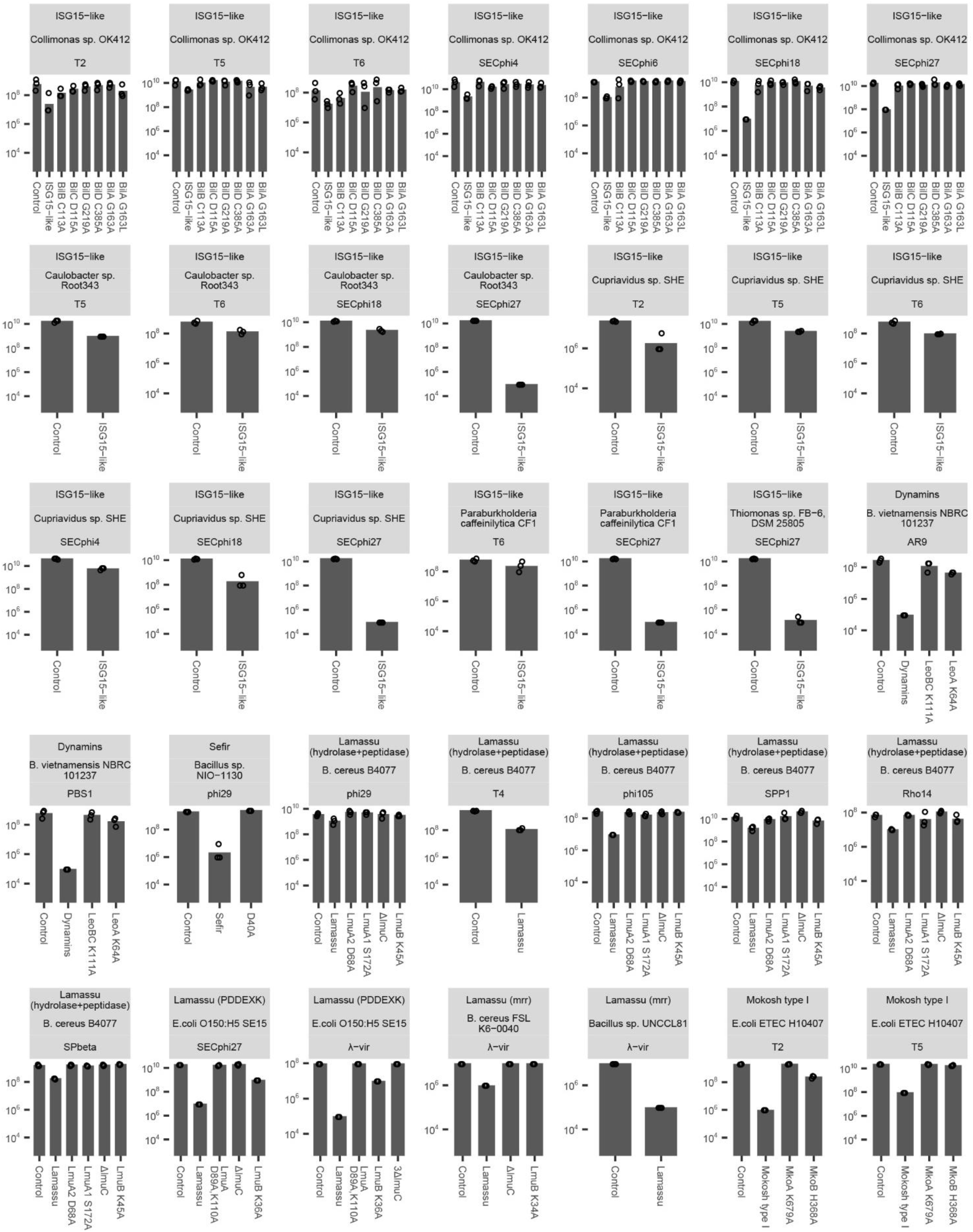
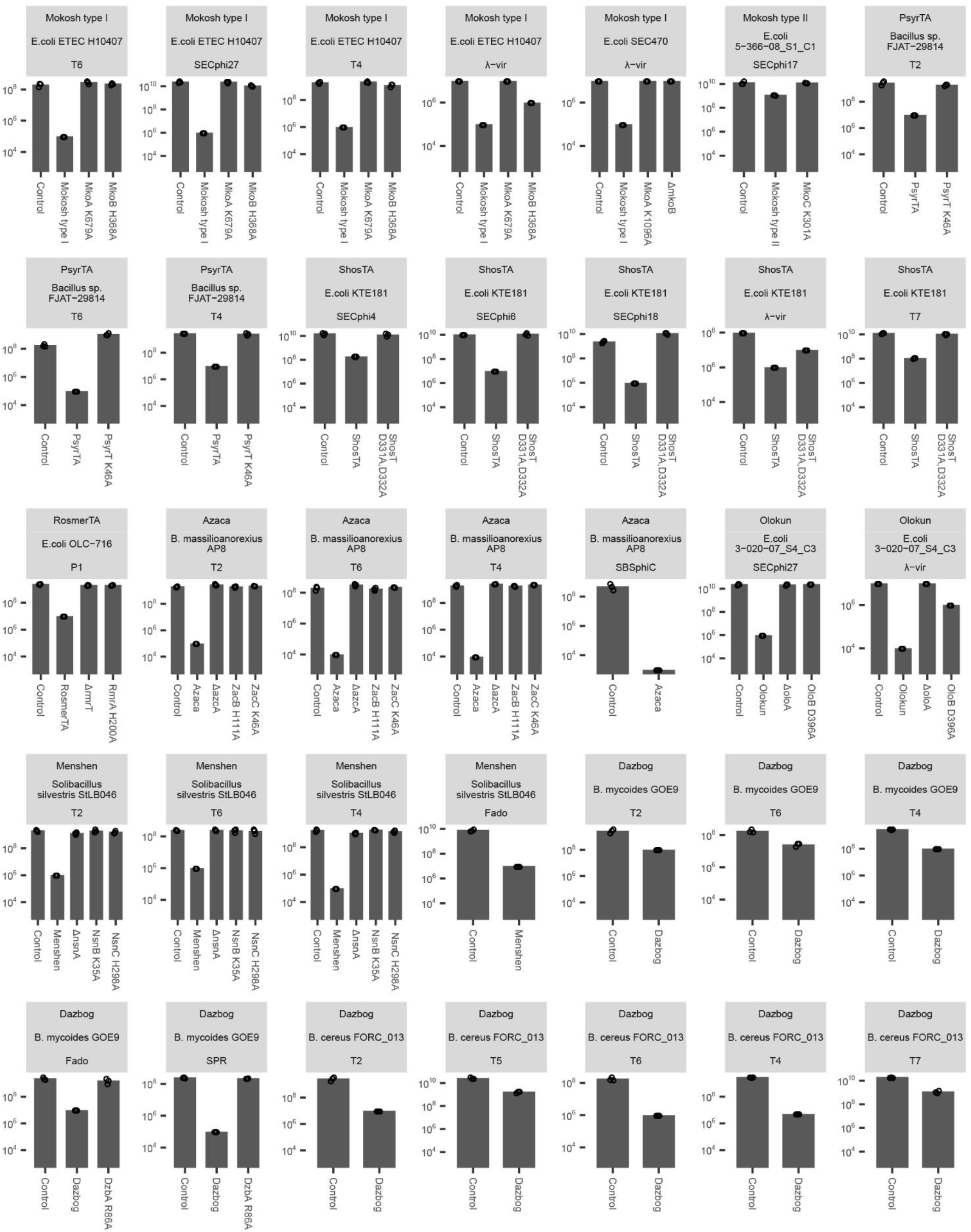
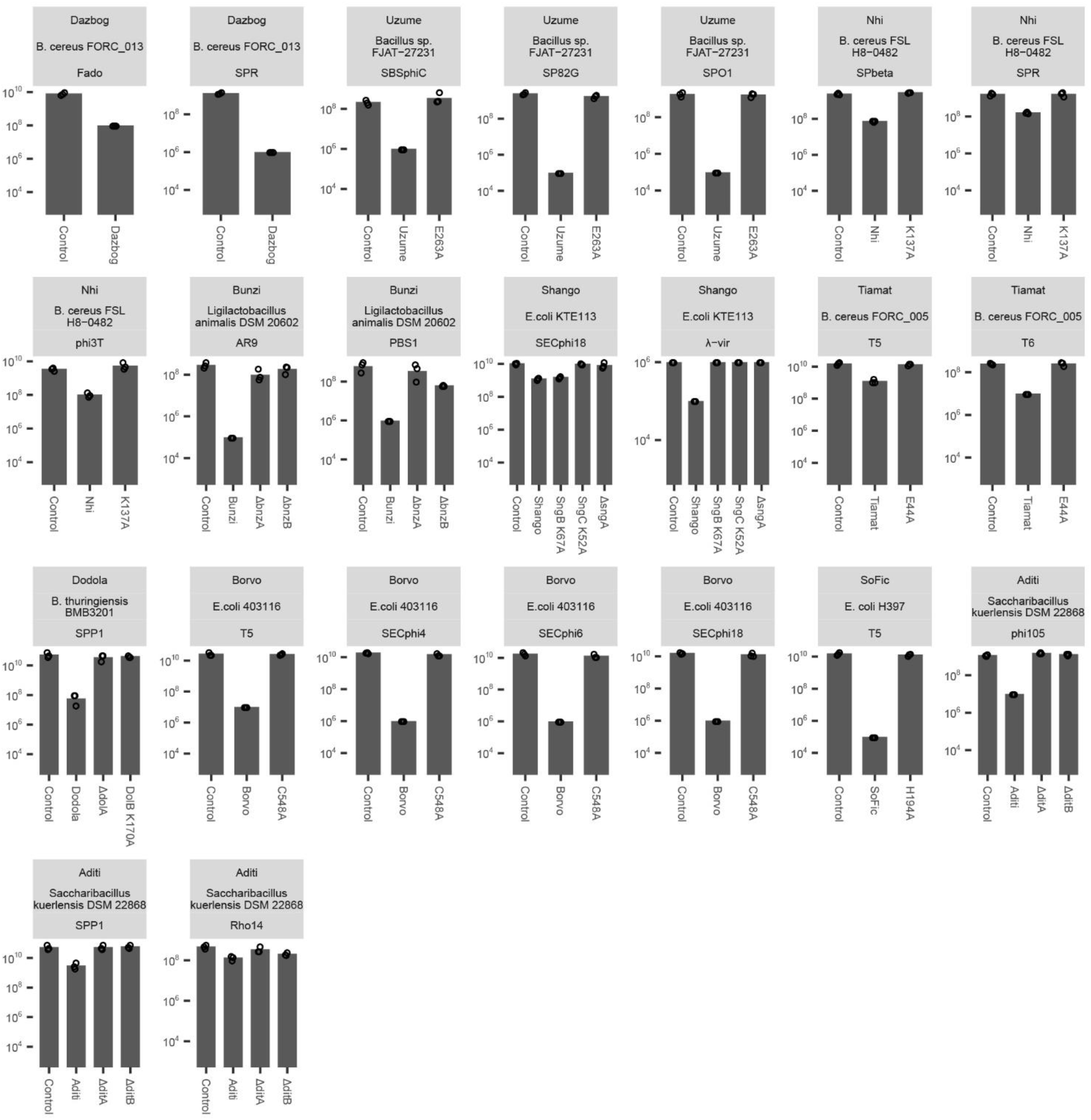
Plaque assay results for all defensive systems described in this study. Efficiency of plating (EOP) of phages infecting *E. coli* or *B. subtilis* with and without WT and mutated systems. Data shown only for phages for which defense was detected. Data represent plaque-forming units (PFU) per ml; bar graphs represent average of three replicates, with individual data points overlaid.

**Figure S2.**
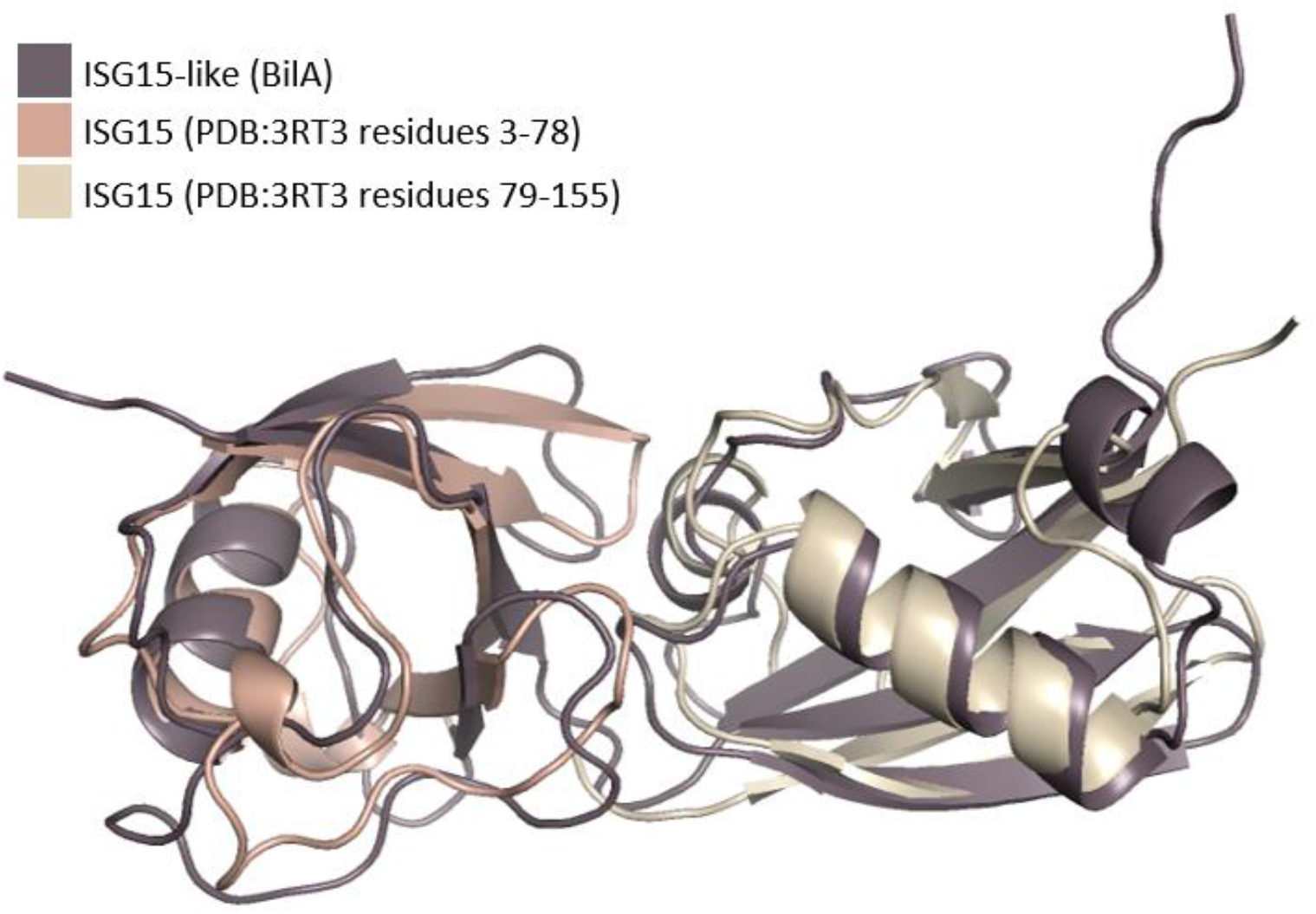
Structural alignment of the ISG15-like protein BilA from *Collimonas sp*. OK412 and the human ISG15. (PDB:3RT3 chain B). Each ubiquitin-like domain from ISG15 was aligned separately. Alignment was performed using PDBeFold (Krissinel and Henrick, 2004) and visualized using PyMOL (Schrödinger, 2015).

**Figure S3.**
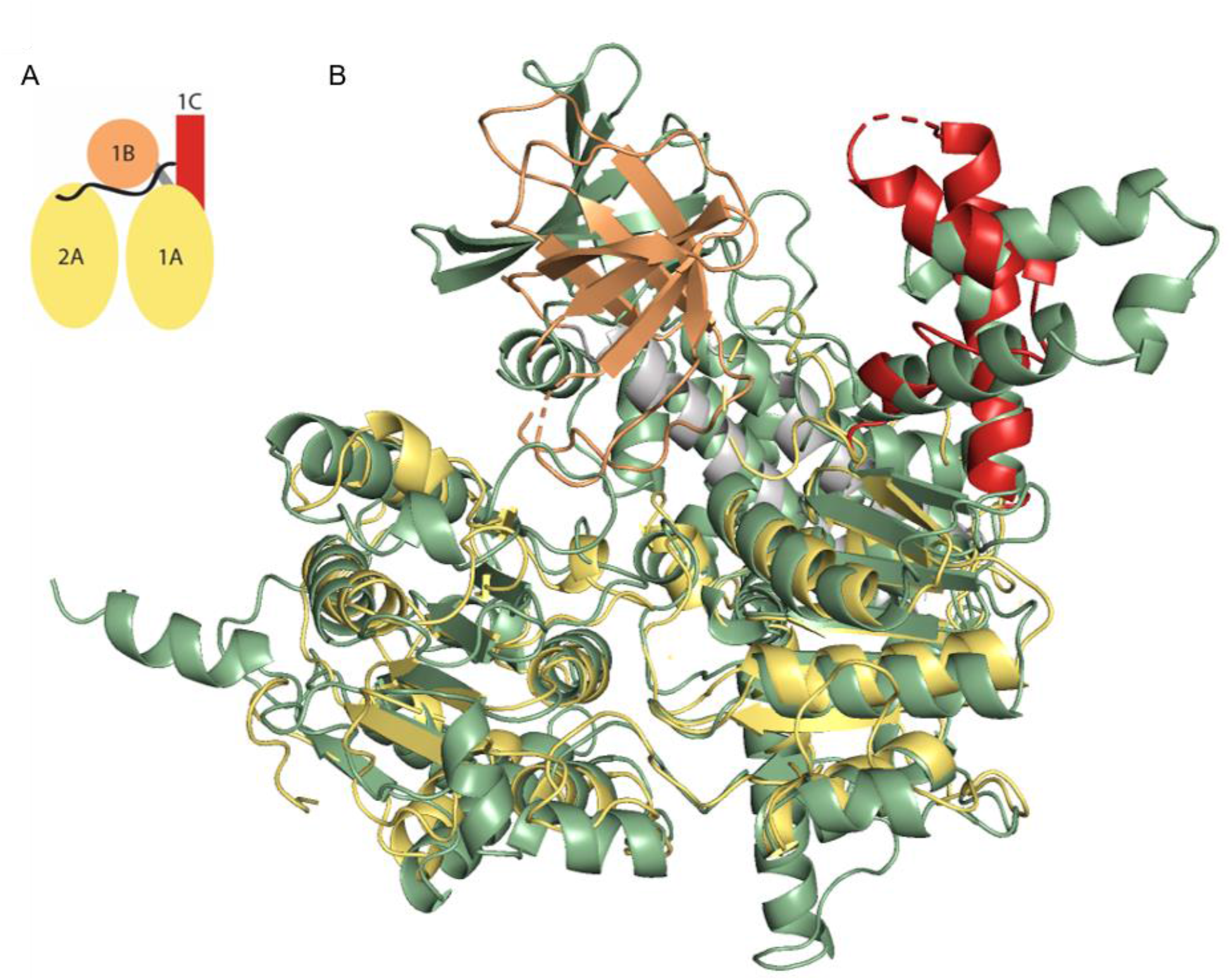
Structural homology of Mokosh protein MkoA to the human Upf1 helicase. (A) Schematic representation of Upf1 family helicases. The Upf1 protein has two RecA domains (yellow) and two additional domains (orange and red). Domains are colored as in (Chakrabarti et al., 2011; Gowravaram et al., 2018). (B) Structural alignment of the AlphaFold-predicted structure of the helicase domain (residues 460-1215) of Mokosh MkoA protein from *E. coli* ETEC H10407 (green) and Upf1 (PDB:6EJ5).

**Figure S4.**
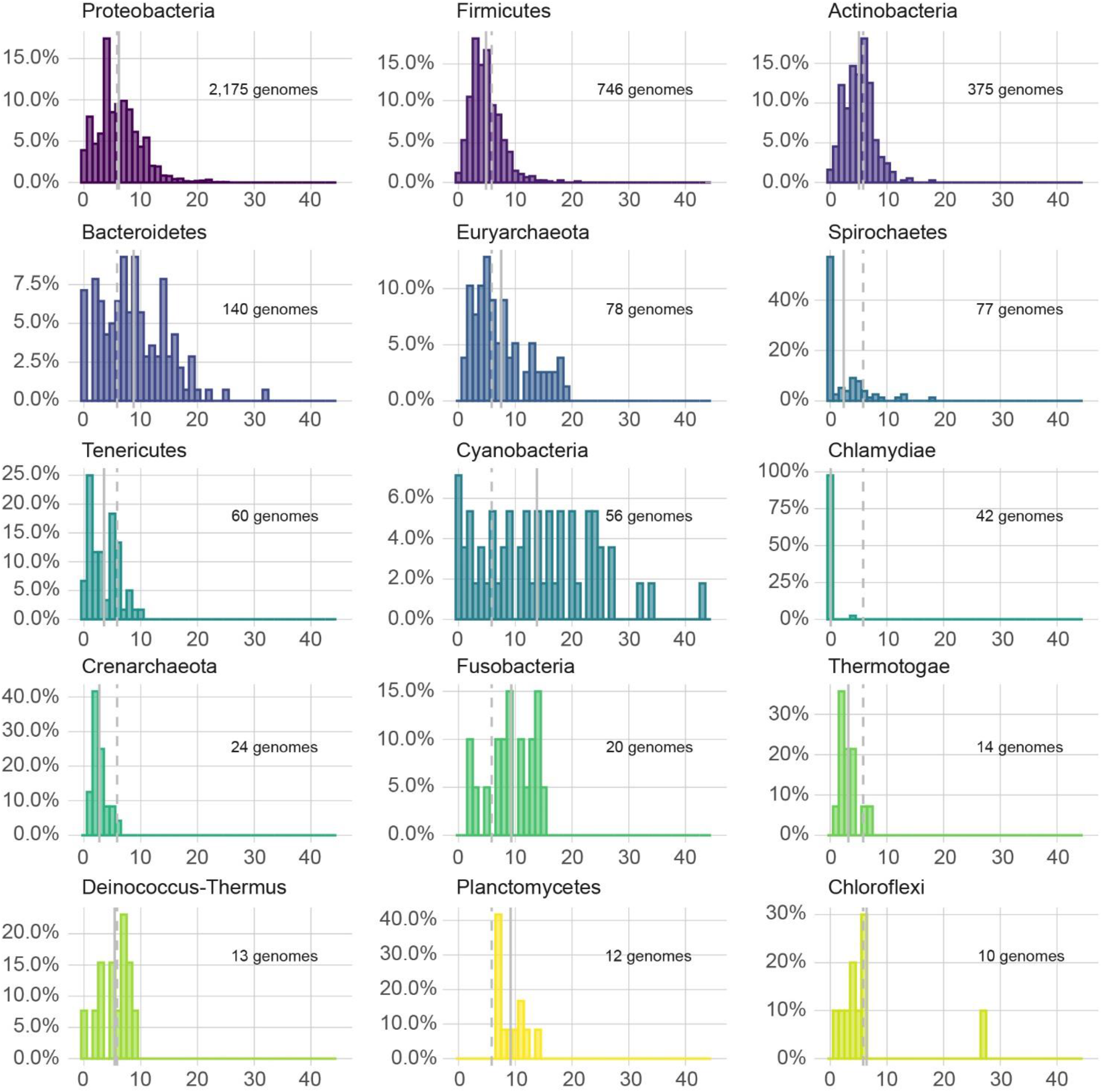
Phylum-specific distribution of defense systems. X-axis represents the number of defense systems per genome; Y-axis represents the fraction of genomes within the phylum. Dashed line is the average out of the entire set of 3,895 genomes; Solid gray line is the average within the phylum. Only phyla with at least 10 finished genomes in our database are presented. Some of the data presented here are also presented in Figure 5.

## References

Alekhina, O., Valkovicova, L., and Turna, J. (2011). Study of membrane attachment and in vivo colocalization of TerB protein from uropathogenic Escherichia coli KL53. Gen. Physiol. Biophys. 30, 286–292.

Anantharaman, V., Iyer, L.M., and Aravind, L. (2012). Ter-dependent stress response systems: novel pathways related to metal sensing, production of a nucleoside-like metabolite, and DNA-processing. Mol. Biosyst. 8, 3142–3165.

Aziz, R.K., Bartels, D., Best, A.A., DeJongh, M., Disz, T., Edwards, R.A., Formsma, K., Gerdes, S., Glass, E.M., Kubal, M., et al. (2008). The RAST Server: rapid annotations using subsystems technology. BMC Genomics 9, 75.

Bankevich, A., Nurk, S., Antipov, D., Gurevich, A.A., Dvorkin, M., Kulikov, A.S., Lesin, V.M., Nikolenko, S.I., Pham, S., Prjibelski, A.D., et al. (2012). SPAdes: a new genome assembly algorithm and its applications to single-cell sequencing. J. Comput. Biol. 19, 455–477.

Bari, S.M.N., Chou-Zheng, L., Howell, O., Hossain, M., Hill, C.M., Boyle, T.A., Cater, K., Dandu, V.S., Thomas, A., Aslan, B., et al. (2022). A unique mode of nucleic acid immunity performed by a multifunctional bacterial enzyme. Cell Host Microbe 30, 570-582.e7.

Baym, M., Kryazhimskiy, S., Lieberman, T.D., Chung, H., Desai, M.M., and Kishony, R. (2015). Inexpensive multiplexed library preparation for megabase-sized genomes. PLoS One 10, e0128036.

Berman, H.M., Westbrook, J., Feng, Z., Gilliland, G., Bhat, T.N., Weissig, H., Shindyalov, I.N., and Bourne, P.E. (2000). The Protein Data Bank. Nucleic Acids Res. 28, 235–242.

Bernheim, A., and Sorek, R. (2020). The pan-immune system of bacteria: antiviral defence as a community resource. Nat. Rev. Microbiol. 18, 113–119.

Bernheim, A., Millman, A., Ofir, G., Meitav, G., Avraham, C., Shomar, H., Rosenberg, M.M., Tal, N., Melamed, S., Amitai, G., et al. (2021). Prokaryotic viperins produce diverse antiviral molecules. Nature 589, 120–124.

Blower, T.R., Pei, X.Y., Short, F.L., Fineran, P.C., Humphreys, D.P., Luisi, B.F., and Salmond, G.P.C. (2011). A processed noncoding RNA regulates an altruistic bacterial antiviral system. Nat. Struct. Mol. Biol. 18, 185–190.

Bobonis, J., Mateus, A., Pfalz, B., Garcia-Santamarina, S., Galardini, M., Kobayashi, C., Stein, F., Savitski, M.M., Elfenbein, J.R., Andrews-Poymenis, H., et al. (2020). Bacterial retrons encode tripartite toxin/antitoxin systems. BioRxiv 2020.06.22.160168.

Chakrabarti, S., Jayachandran, U., Bonneau, F., Fiorini, F., Basquin, C., Domcke, S., Le Hir, H., and Conti, E. (2011). Molecular mechanisms for the RNA-dependent ATPase activity of Upf1 and its regulation by Upf2. Mol. Cell 41, 693–703.

Chen, I.M.A., Chu, K., Palaniappan, K., Pillay, M., Ratner, A., Huang, J., Huntemann, M., Varghese, N., White, J.R., Seshadri, R., et al. (2019). IMG/M v.5.0: An integrated data management and comparative analysis system for microbial genomes and microbiomes. Nucleic Acids Res. 47, D666–D677.

Cohen, D., Melamed, S., Millman, A., Shulman, G., Oppenheimer-Shaanan, Y., Kacen, A., Doron, S., Amitai, G., and Sorek, R. (2019). Cyclic GMP-AMP signalling protects bacteria against viral infection. Nature 574, 691–695.

Crameri, M., Bauer, M., Caduff, N., Walker, R., Steiner, F., Franzoso, F.D., Gujer, C., Boucke, K., Kucera, T., Zbinden, A., et al. (2018). MxB is an interferon-induced restriction factor of human herpesviruses. Nat. Commun. 9, 1980.

D’Cunha, J., Knight, E., Haas, A.L., Truitt, R.L., and Borden, E.C. (1996). Immunoregulatory properties of ISG15, an interferon-induced cytokine. Proc. Natl. Acad. Sci. U. S. A. 93, 211–215.

Dastur, A., Beaudenon, S., Kelley, M., Krug, R.M., and Huibregtse, J.M. (2006). Herc5, an interferon-induced HECT E3 enzyme, is required for conjugation of ISG15 in human cells. J. Biol. Chem. 281, 4334–4338.

Doron, S., Melamed, S., Ofir, G., Leavitt, A., Lopatina, A., Keren, M., Amitai, G., and Sorek, R. (2018). Systematic discovery of antiphage defense systems in the microbial pangenome. Science 359, eaar4120.

Dzimianski, J. V, Scholte, F.E.M., Bergeron, É., and Pegan, S.D. (2019). ISG15: It’s Complicated. J. Mol. Biol. 431, 4203–4216.

Eddy, S.R. (2011). Accelerated Profile HMM Searches. PLoS Comput. Biol. 7, e1002195.

El-Gebali, S., Mistry, J., Bateman, A., Eddy, S.R., Luciani, A., Potter, S.C., Qureshi, M., Richardson, L.J., Salazar, G.A., Smart, A., et al. (2019). The Pfam protein families database in 2019. Nucleic Acids Res. 47, D427–D432.

Engel, P., Goepfert, A., Stanger, F. V, Harms, A., Schmidt, A., Schirmer, T., and Dehio, C. (2012). Adenylylation control by intraor intermolecular active-site obstruction in Fic proteins. Nature 482, 107–110.

Fleckenstein, J.M., Lindler, L.E., Elsinghorst, E.A., and Dale, J.B. (2000). Identification of a gene within a pathogenicity island of enterotoxigenic Escherichia coli H10407 required for maximal secretion of the heat-labile enterotoxin. Infect. Immun. 68, 2766–2774.

Freitas, B.T., Scholte, F.E.M., Bergeron, É., and Pegan, S.D. (2020). How ISG15 combats viral infection. Virus Res. 286, 198036.

Frese, M., Kochs, G., Feldmann, H., Hertkorn, C., and Haller, O. (1996). Inhibition of bunyaviruses, phleboviruses, and hantaviruses by human MxA protein. J. Virol. 70, 915–923.

Gao, L., Altae-Tran, H., Böhning, F., Makarova, K.S., Segel, M., Schmid-Burgk, J.L., Koob, J., Wolf, Y.I., Koonin, E. V, and Zhang, F. (2020). Diverse enzymatic activities mediate antiviral immunity in prokaryotes. Science 369, 1077–1084.

Garb, J., Lopatina, A., Bernheim, A., Zaremba, M., Siksnys, V., Melamed, S., Leavitt, A., Millman, A., Amitai, G., and Sorek, R. (2021). Multiple phage resistance systems inhibit infection via SIR2-dependent NAD+ depletion. BioRxiv 2021.12.14.472415.

Goldfarb, T., Sberro, H., Weinstock, E., Cohen, O., Doron, S., Charpak-Amikam, Y., Afik, S., Ofir, G., and Sorek, R. (2015). BREX is a novel phage resistance system widespread in microbial genomes. EMBO J. 34, 169–183.

Gowravaram, M., Bonneau, F., Kanaan, J., Maciej, V.D., Fiorini, F., Raj, S., Croquette, V., Le Hir, H., and Chakrabarti, S. (2018). A conserved structural element in the RNA helicase UPF1 regulates its catalytic activity in an isoform-specific manner. Nucleic Acids Res. 46, 2648–2659.

Guo, L., Sattler, L., Shafqat, S., Graumann, P.L., and Bramkamp, M. (2022). A Bacterial Dynamin-Like Protein Confers a Novel Phage Resistance Strategy on the Population Level in Bacillus subtilis. MBio e0375321.

Haering, C.H., Löwe, J., Hochwagen, A., and Nasmyth, K. (2002). Molecular architecture of SMC proteins and the yeast cohesin complex. Mol. Cell 9, 773–788.

Haller, O., and Kochs, G. (2020). Mx genes: host determinants controlling influenza virus infection and trans-species transmission. Hum. Genet. 139, 695–705.

Haller, O., Staeheli, P., Schwemmle, M., and Kochs, G. (2015). Mx GTPases: dynamin-like antiviral machines of innate immunity. Trends Microbiol. 23, 154–163.

Hampton, H.G., Watson, B.N.J., and Fineran, P.C. (2020). The arms race between bacteria and their phage foes. Nature 577, 327–336.

Harms, A., Stanger, F. V, and Dehio, C. (2016). Biological Diversity and Molecular Plasticity of FIC Domain Proteins. Annu. Rev. Microbiol. 70, 341–360.

Jaskólska, M., Adams, D.W., and Blokesch, M. (2022). Two defence systems eliminate plasmids from seventh pandemic Vibrio cholerae. Nature 604, 323–329.

Johnson, A.G., Wein, T., Mayer, M.L., Duncan-Lowey, B., Yirmiya, E., Oppenheimer-Shaanan, Y., Amitai, G., Sorek, R., and Kranzusch, P.J. (2022). Bacterial gasdermins reveal an ancient mechanism of cell death. Science 375, 221–225.

Jumper, J., Evans, R., Pritzel, A., Green, T., Figurnov, M., Ronneberger, O., Tunyasuvunakool, K., Bates, R., Žídek, A., Potapenko, A., et al. (2021). Highly accurate protein structure prediction with AlphaFold. Nature 596, 583–589.

Käshammer, L., Saathoff, J.-H., Lammens, K., Gut, F., Bartho, J., Alt, A., Kessler, B., and Hopfner, K.-P. (2019). Mechanism of DNA End Sensing and Processing by the Mre11-Rad50 Complex. Mol. Cell 76, 382-394.e6.

Katoh, K., Misawa, K., Kuma, K., and Miyata, T. (2002). MAFFT: a novel method for rapid multiple sequence alignment based on fast Fourier transform. Nucleic Acids Res. 30, 3059–3066.

Kever, L., Hardy, A., Luthe, T., Hünnefeld, M., Gätgens, C., Milke, L., Wiechert, J., Wittmann, J., Moraru, C., Marienhagen, J., et al. (2021). Aminoglycoside antibiotics inhibit phage infection by blocking an early step of the phage infection cycle. BioRxiv 2021.05.02.442312.

Kim, Y.K., Furic, L., Desgroseillers, L., and Maquat, L.E. (2005). Mammalian Staufen1 recruits Upf1 to specific mRNA 3’UTRs so as to elicit mRNA decay. Cell 120, 195–208.

Kimelman, A., Levy, A., Sberro, H., Kidron, S., Leavitt, A., Amitai, G., Yoder-Himes, D.R., Wurtzel, O., Zhu, Y., Rubin, E.M., et al. (2012). A vast collection of microbial genes that are toxic to bacteria. Genome Res. 22, 802–809.

Koga, M., Otsuka, Y., Lemire, S., and Yonesaki, T. (2011). Escherichia coli rnlA and rnlB compose a novel toxin-antitoxin system. Genetics 187, 123–130.

Koopal, B., Potocnik, A., Mutte, S.K., Aparicio-Maldonado, C., Lindhoud, S., Vervoort, J.J.M., Brouns, S.J.J., and Swarts, D.C. (2022). Short prokaryotic Argonaute systems trigger cell death upon detection of invading DNA. Cell 185, 1471-1486.e19.

Krishnan, A., Burroughs, A.M., Iyer, L.M., and Aravind, L. (2020). Comprehensive classification of ABC ATPases and their functional radiation in nucleoprotein dynamics and biological conflict systems. Nucleic Acids Res. 48, 10045–10075.

Krissinel, E., and Henrick, K. (2004). Secondary-structure matching (SSM), a new tool for fast protein structure alignment in three dimensions. Acta Crystallogr. D. Biol. Crystallogr. 60, 2256–2268.

Kronheim, S., Daniel-Ivad, M., Duan, Z., Hwang, S., Wong, A.I., Mantel, I., Nodwell, J.R., and Maxwell, K.L. (2018). A chemical defence against phage infection. Nature 564, 283–286.

Krug, R.M., Shaw, M., Broni, B., Shapiro, G., and Haller, O. (1985). Inhibition of influenza viral mRNA synthesis in cells expressing the interferon-induced Mx gene product. J. Virol. 56, 201–206.

Kuzmenko, A., Oguienko, A., Esyunina, D., Yudin, D., Petrova, M., Kudinova, A., Maslova, O., Ninova, M., Ryazansky, S., Leach, D., et al. (2020). DNA targeting and interference by a bacterial Argonaute nuclease. Nature 587, 632–637.

Ledvina, H.E., Ye, Q., Gu, Y., Quan, Y., Lau, R.K., Zhou, H., Corbett, K.D., and Whiteley, A.T. (2022). cGASylation by a bacterial E1-E2 fusion protein primes antiviral immune signaling. BioRxiv 2022.03.31.486616.

LeRoux, M., and Laub, M.T. (2022). Toxin-Antitoxin Systems as Phage Defense Elements. Annu. Rev. Microbiol. 10.1146/annurev-micro-020722–013730.

LeRoux, M., Srikant, S., Littlehale, M.H., Teodoro, G., Doron, S., Badiee, M., Leung, A.K.L., Sorek, R., and Laub, M.T. (2021). The DarTG toxin-antitoxin system provides phage defense by ADP-ribosylating viral DNA. BioRxiv 2021.09.27.462013.

Lopatina, A., Tal, N., and Sorek, R. (2020). Abortive Infection: Bacterial Suicide as an Antiviral Immune Strategy. Annu. Rev. Virol. 7, 371–384.

Losada, A., Hirano, M., and Hirano, T. (1998). Identification of Xenopus SMC protein complexes required for sister chromatid cohesion. Genes Dev. 12, 1986–1997.

Lowey, B., Whiteley, A.T., Keszei, A.F.A., Morehouse, B.R., Mathews, I.T., Antine, S.P., Cabrera, V.J., Kashin, D., Niemann, P., Jain, M., et al. (2020). CBASS Immunity Uses CARF-Related Effectors to Sense 3’-5’- and 2’-5’-Linked Cyclic Oligonucleotide Signals and Protect Bacteria from Phage Infection. Cell 182, 38-49.e17.

Luo, P., He, X., Liu, Q., and Hu, C. (2015). Developing Universal Genetic Tools for Rapid and Efficient Deletion Mutation in Vibrio Species Based on Suicide T-Vectors Carrying a Novel Counterselectable Marker, vmi480. PLoS One 10, e0144465.

Makarova, K.S., Wolf, Y.I., Snir, S., and Koonin, E. V. (2011). Defense Islands in Bacterial and Archaeal Genomes and Prediction of Novel Defense Systems. J. Bacteriol. 193, 6039–6056.

Makarova, K.S., Wolf, Y.I., Iranzo, J., Shmakov, S.A., Alkhnbashi, O.S., Brouns, S.J.J., Charpentier, E., Cheng, D., Haft, D.H., Horvath, P., et al. (2020). Evolutionary classification of CRISPR-Cas systems: a burst of class 2 and derived variants. Nat. Rev. Microbiol. 18, 67–83.

Matelska, D., Steczkiewicz, K., and Ginalski, K. (2017). Comprehensive classification of the PIN domain-like superfamily. Nucleic Acids Res. 45, 6995–7020.

Mazzocco, A., Waddell, T.E., Lingohr, E., and Johnson, R.P. (2009). Enumeration of bacteriophages using the small drop plaque assay system. Methods Mol. Biol. 501, 81–85.

Mesev, E. V, LeDesma, R.A., and Ploss, A. (2019). Decoding type I and III interferon signalling during viral infection. Nat. Microbiol. 4, 914–924.

Mestre, M.R., González-Delgado, A., Gutiérrez-Rus, L.I., Martínez-Abarca, F., and Toro, N. (2020). Systematic prediction of genes functionally associated with bacterial retrons and classification of the encoded tripartite systems. Nucleic Acids Res. 48, 12632–12647.

Michaelis, C., Ciosk, R., and Nasmyth, K. (1997). Cohesins: chromosomal proteins that prevent premature separation of sister chromatids. Cell 91, 35–45.

Michie, K.A., Boysen, A., Low, H.H., Møller-Jensen, J., and Löwe, J. (2014). LeoA, B and C from enterotoxigenic Escherichia coli (ETEC) are bacterial dynamins. PLoS One 9, e107211.

Millman, A., Bernheim, A., Stokar-Avihail, A., Fedorenko, T., Voichek, M., Leavitt, A., Oppenheimer-Shaanan, Y., and Sorek, R. (2020a). Bacterial Retrons Function In Anti-Phage Defense. Cell 183, 1551-1561.e12.

Millman, A., Melamed, S., Amitai, G., and Sorek, R. (2020b). Diversity and classification of cyclic-oligonucleotide-based anti-phage signalling systems. Nat. Microbiol. 5, 1608–1615.

Mirdita, M., von den Driesch, L., Galiez, C., Martin, M.J., Söding, J., and Steinegger, M. (2017). Uniclust databases of clustered and deeply annotated protein sequences and alignments. Nucleic Acids Res. 45, D170–D176.

Morehouse, B.R., Govande, A.A., Millman, A., Keszei, A.F.A., Lowey, B., Ofir, G., Shao, S., Sorek, R., and Kranzusch, P.J. (2020). STING cyclic dinucleotide sensing originated in bacteria. Nature 586, 429–433.

Mortier-Barrière, I., Velten, M., Dupaigne, P., Mirouze, N., Piétrement, O., McGovern, S., Fichant, G., Martin, B., Noirot, P., Le Cam, E., et al. (2007). A key presynaptic role in transformation for a widespread bacterial protein: DprA conveys incoming ssDNA to RecA. Cell 130, 824–836.

Narasimhan, J., Wang, M., Fu, Z., Klein, J.M., Haas, A.L., and Kim, J.-J.P. (2005). Crystal structure of the interferon-induced ubiquitin-like protein ISG15. J. Biol. Chem. 280, 27356–27365.

Novatchkova, M., Leibbrandt, A., Werzowa, J., Neubüser, A., and Eisenhaber, F. (2003). The STIR-domain superfamily in signal transduction, development and immunity. Trends Biochem. Sci. 28, 226–229.

Ofir, G., and Sorek, R. (2018). Contemporary Phage Biology: From Classic Models to New Insights. Cell 172, 1260–1270.

Ofir, G., Herbst, E., Baroz, M., Cohen, D., Millman, A., Doron, S., Tal, N., Malheiro, D.B.A., Malitsky, S., Amitai, G., et al. (2021). Antiviral activity of bacterial TIR domains via immune signalling molecules. Nature 600, 116–120.

Oliveira, P.H., Touchon, M., and Rocha, E.P.C. (2014). The interplay of restriction-modification systems with mobile genetic elements and their prokaryotic hosts. Nucleic Acids Res. 42, 10618–10631.

Onishi, R.M., Park, S.J., Hanel, W., Ho, A.W., Maitra, A., and Gaffen, S.L. (2010). SEF/IL-17R (SEFIR) is not enough: an extended SEFIR domain is required for il-17RA-mediated signal transduction. J. Biol. Chem. 285, 32751–32759.

Payne, L.J., Todeschini, T.C., Wu, Y., Perry, B.J., Ronson, C.W., Fineran, P.C., Nobrega, F.L., and Jackson, S.A. (2021). Identification and classification of antiviral defence systems in bacteria and archaea with PADLOC reveals new system types. Nucleic Acids Res. 49, 10868–10878.

Perng, Y.-C., and Lenschow, D.J. (2018). ISG15 in antiviral immunity and beyond. Nat. Rev. Microbiol. 16, 423–439.

Potts, P.R., Porteus, M.H., and Yu, H. (2006). Human SMC5/6 complex promotes sister chromatid homologous recombination by recruiting the SMC1/3 cohesin complex to double-strand breaks. EMBO J. 25, 3377–3388.

Ramachandran, R., and Schmid, S.L. (2018). The dynamin superfamily. Curr. Biol. 28, R411–R416.

Rousset, F., Cabezas-Caballero, J., Piastra-Facon, F., Fernández-Rodríguez, J., Clermont, O., Denamur, E., Rocha, E.P.C., and Bikard, D. (2021). The impact of genetic diversity on gene essentiality within the Escherichia coli species. Nat. Microbiol. 6, 301–312.

Rousset, F., Depardieu, F., Miele, S., Dowding, J., Laval, A.-L., Lieberman, E., Garry, D., Rocha, E.P.C., Bernheim, A., and Bikard, D. (2022). Phages and their satellites encode hotspots of antiviral systems. Cell Host Microbe 10.1016/j.chom.2022.02.018.

Russell, C.W., and Mulvey, M.A. (2015). The Extraintestinal Pathogenic Escherichia coli Factor RqlI Constrains the Genotoxic Effects of the RecQ-Like Helicase RqlH. PLoS Pathog. 11, e1005317.

Ryzhakov, G., Blazek, K., and Udalova, I.A. (2011). Evolution of vertebrate immunity: sequence and functional analysis of the SEFIR domain family member Act1. J. Mol. Evol. 72, 521–530.

Sberro, H., Leavitt, A., Kiro, R., Koh, E., Peleg, Y., Qimron, U., and Sorek, R. (2013). Discovery of functional toxin/antitoxin systems in bacteria by shotgun cloning. Mol. Cell 50, 136–148.

Schrödinger, L. (2015). The PyMOL Molecular Graphics System, Version 2.5.2.

Serrero, M.C., Girault, V., Weigang, S., Greco, T.M., Ramos-Nascimento, A., Anderson, F., Piras, A., Martinez, A.H., Hertzog, J., Binz, A., et al. (2022). The interferon-inducible GTPase MxB promotes capsid disassembly and genome release of herpesviruses. BioRxiv 2022.01.25.477704.

Singh, M., Bindal, G., Misra, C.S., and Rath, D. (2022). The era of Cas12 and Cas13 CRISPR-based disease diagnosis. Crit. Rev. Microbiol. 1–16.

Steinegger, M., and Söding, J. (2017). MMseqs2 enables sensitive protein sequence searching for the analysis of massive data sets. Nat. Biotechnol. 35, 1026–1028.

Steinegger, M., Meier, M., Mirdita, M., Vöhringer, H., Haunsberger, S.J., and Söding, J. (2019). HH-suite3 for fast remote homology detection and deep protein annotation. BMC Bioinformatics 20, 473.

Storey, J.D., Bass, A.J., Dabney, A., and Robinson, D. (2019). qvalue: Q-value estimation for false discovery rate control. http://github.com/jdstorey/qvalue.

Tal, N., and Sorek, R. (2022). SnapShot: Bacterial immunity. Cell 185, 578-578.e1.

Tal, N., Millman, A., Stokar-Avihail, A., Fedorenko, T., Leavitt, A., Melamed, S., Yirmiya, E., Avraham, C., Amitai, G., and Sorek, R. (2021a). Antiviral defense via nucleotide depletion in bacteria. BioRxiv 2021.04.26.441389.

Tal, N., Morehouse, B.R., Millman, A., Stokar-Avihail, A., Avraham, C., Fedorenko, T., Yirmiya, E., Herbst, E., Brandis, A., Mehlman, T., et al. (2021b). Cyclic CMP and cyclic UMP mediate bacterial immunity against phages. Cell 184, 5728-5739.e16.

Tesson, F., Hervé, A., Touchon, M., d’Humières, C., Cury, J., and Bernheim, A. (2021). Systematic and quantitative view of the antiviral arsenal of prokaryotes. BioRxiv 2021.09.02.458658.

Wein, T., and Sorek, R. (2022). Bacterial origins of human cell-autonomous innate immune mechanisms. Nat. Rev. Immunol. 10.1038/s41577-022-00705–4.

Whelan, K.F., Sherburne, R.K., and Taylor, D.E. (1997). Characterization of a region of the IncHI2 plasmid R478 which protects Escherichia coli from toxic effects specified by components of the tellurite, phage, and colicin resistance cluster. J. Bacteriol. 179, 63–71.

Whiteley, A.T., Eaglesham, J.B., de Oliveira Mann, C.C., Morehouse, B.R., Lowey, B., Nieminen, E.A., Danilchanka, O., King, D.S., Lee, A.S.Y., Mekalanos, J.J., et al. (2019). Bacterial cGAS-like enzymes synthesize diverse nucleotide signals. Nature 567, 194–199.

Wilson, G.A., and Bott, K.F. (1968). Nutritional factors influencing the development of competence in the Bacillus subtilis transformation system. J. Bacteriol. 95, 1439–1449.

Wu, B., Gong, J., Liu, L., Li, T., Wei, T., and Bai, Z. (2012). Evolution of prokaryotic homologues of the eukaryotic SEFIR protein domain. Gene 492, 160–166.

Yang, H., Zhu, Y., Chen, X., Li, X., Ye, S., and Zhang, R. (2018). Structure of a prokaryotic SEFIR domain reveals two novel SEFIR-SEFIR interaction modes. J. Struct. Biol. 203, 81–89.

Ye, Q., Lau, R.K., Mathews, I.T., Birkholz, E.A., Watrous, J.D., Azimi, C.S., Pogliano, J., Jain, M., and Corbett, K.D. (2020). HORMA Domain Proteins and a Trip13-like ATPase Regulate Bacterial cGAS-like Enzymes to Mediate Bacteriophage Immunity. Mol. Cell 77, 709-722.e7.

Zaremba, M., Dakineviciene, D., Golovinas, E., Stankunas, E., Lopatina, A., Sorek, R., Manakova, E., Ruksenaite, A., Silanskas, A., Asmontas, S., et al. (2021). Sir2-domain associated short prokaryotic Argonautes provide defence against invading mobile genetic elements through NAD+ depletion. BioRxiv 2021.12.14.472599.

Zhang, B., Liu, C., Qian, W., Han, Y., Li, X., and Deng, J. (2014). Structure of the unique SEFIR domain from human interleukin 17 receptor A reveals a composite ligand-binding site containing a conserved α-helix for Act1 binding and IL-17 signaling. Acta Crystallogr. D. Biol. Crystallogr. 70, 1476–1483.

Zhao, C., Beaudenon, S.L., Kelley, M.L., Waddell, M.B., Yuan, W., Schulman, B.A., Huibregtse, J.M., and Krug, R.M. (2004). The UbcH8 ubiquitin E2 enzyme is also the E2 enzyme for ISG15, an IFN-alpha/beta-induced ubiquitin-like protein. Proc. Natl. Acad. Sci. U. S. A. 101, 7578–7582.

Zimmermann, L., Stephens, A., Nam, S.-Z., Rau, D., Kübler, J., Lozajic, M., Gabler, F., Söding, J., Lupas, A.N., and Alva, V. (2018). A Completely Reimplemented MPI Bioinformatics Toolkit with a New HHpred Server at its Core. J. Mol. Biol. 430, 2237–2243.

